# Translation inhibition from a distance: the small RNA SgrS interferes with a ribosomal protein S1-dependent enhancer

**DOI:** 10.1101/808485

**Authors:** Muhammad S. Azam, Carin K. Vanderpool

## Abstract

Many bacterial small RNAs (sRNAs) efficiently inhibit translation of target mRNAs by forming a duplex that sequesters the Shine-Dalgarno (SD) sequence or start codon and prevents formation of the translation initiation complex. There are a growing number of examples of sRNA-mRNA binding interactions distant from the SD region, but how these mediate translational regulation remains unclear. Our previous work in *Escherichia coli* and *Salmonella* identified a mechanism of translational repression of *manY* mRNA by the sRNA SgrS through a binding interaction upstream of the *manY* SD. Here, we report that SgrS forms a duplex with a uridine-rich translation-enhancing element in the *manY* 5’ untranslated region. Notably, we show that the enhancer is ribosome-dependent and that the small ribosomal subunit protein S1 interacts with the enhancer to promote translation of *manY*. In collaboration with the chaperone protein Hfq, SgrS interferes with the interaction between the translation enhancer and ribosomal protein S1 to repress translation of *manY* mRNA. Since bacterial translation is often modulated by enhancer-like elements upstream of the SD, sRNA-mediated enhancer silencing could be a common mode of gene regulation.

## Introduction

Microbes respond to changes in environmental conditions using a variety of mechanisms to modify gene expression resulting in changes in cell structure and function. Small RNAs (sRNAs) are found in microbes across the tree of life and are major regulators responsible for post-transcriptional control of gene regulation. sRNAs regulate target mRNA translation and stability by a variety of molecular mechanisms that depend on sRNA-mRNA duplex formation. Early studies characterizing sRNA-mRNA interactions reported sRNA base pairing with sequences around the Shine-Dalgarno (SD) sequence (Mizuno *et al.*, 1984, Schmidt *et al.*, 1995). In vitro structure probing of sRNA-mRNA duplexes and toeprinting assays probing translation initiation complex formation (Argaman & Altuvia, 2000, Geissmann & Touati, 2004, Møller *et al.*, 2002, Bouvier *et al.*, 2008), have revealed that a common mechanism of sRNA-mediated translational repression is steric occlusion of ribosome binding. However, a growing number of examples of translational repression by sRNAs involve base-pairing interactions far from the SD sequence (Desnoyers & Massé, 2012, Holmqvist *et al.*, 2010, Sharma *et al.*, 2007, Azam & Vanderpool, 2018). Several mechanisms have been reported for regulation of translation via sRNA-mRNA duplex formation distant from ribosome binding sites. The first involves sRNA binding at a site upstream or downstream of the translation start resulting in recruitment of the RNA chaperone Hfq to bind at a site near the SD so that Hfq occludes ribosome binding (Desnoyers & Massé, 2012, Azam & Vanderpool, 2018). Another mechanism involves sRNA sequestration of CA-rich sequences that act as translation enhancer elements (Sharma *et al.*, 2007, Yang *et al.*, 2014). The sRNA GcvB represses translation of several mRNAs using a GU-rich seed region that base pairs with CA-rich target sites located at variable distances upstream of the SD. In the absence of GcvB, the CA-rich sequences enhance mRNA translation by an unknown mechanism. With GcvB present, these enhancer sequences are sequestered and translation of mRNA targets is reduced. A third mechanism of sRNA-mediated translational repression from a distance involves ribosome standby sites. Certain mRNAs with stable secondary structures around the ribosome binding site use upstream ribosome standby sites to promote translation. Ribosome standby sites are located in single-stranded regions where a 30S ribosomal subunit can bind and kinetically compete with RNA folding to access the downstream translation initiation region (Sterk *et al.*, 2018, de Smit & van Duin, 2003, Studer & Joseph, 2006). The toxin-encoding *tisB* mRNA has a ribosome standby site far upstream of the SD, and this site can promote translation of *tisB* only in the absence of an antisense sRNA, IstR, which sequesters the standby site (Romilly *et al.*, 2019, Darfeuille *et al.*, 2007). These examples illustrate the many mechanisms of translational repression mediated by sRNAs binding to mRNAs within and outside the translation initiation region.

The rate-limiting step in protein synthesis is generally considered to be binding of the small ribosomal subunit to the mRNA upstream of the start codon to form the translation initiation complex (Milon *et al.*, 2012, Kozak, 1999). In most cases, the mRNA leader region contains a purine-rich SD sequence (Shine & Dalgarno, 1974) just upstream of the start codon that forms a short duplex with the 3’-end of the 16S rRNA (anti-SD sequence). Some leader regions may contain additional sequence features that can also determine the rate of translation initiation (Miranda-Ríos *et al.*, 2001). Pyrimidine-rich sequences that are sometimes present immediately upstream of the SD can interact with the 30S subunit ribosomal protein S1 and act as translational enhancers (Boni *et al.*, 2001, Boni *et al.*, 1991, Duval *et al.*, 2013). For these mRNAs, formation of the preinitiation complex involves RNA-RNA interactions and RNA-protein interactions that both contribute to the efficiency of translation initiation (Duval *et al.*, 2013, Takahashi *et al.*, 2013, Marzi *et al.*, 2007). sRNAs could interfere with translation initiation by disrupting any of the interactions – RNA-RNA or RNA-protein – that are important for formation of the translation initiation complex.

In this study, we aimed to define the molecular mechanism by which an *E. coli* sRNA, SgrS, regulates translation from a distance. SgrS represses translation of all the cistrons encoded by the *manXYZ* mRNA, which encodes the mannose PTS transporter (Rice & Vanderpool, 2011, Rice *et al.*, 2012). SgrS binds to two distinct sites on *manXYZ* mRNA (Rice *et al.*, 2012, Rice & Vanderpool, 2011, Azam & Vanderpool, 2018). The first site is located within the *manX* coding sequence, and we showed recently that SgrS represses translation of *manX* via a non-canonical mechanism involving SgrS-dependent recruitment of Hfq to a binding site that overlaps the *manX* SD sequence – making SgrS the chaperone-like partner and Hfq the direct repressor of translation (Azam & Vanderpool, 2018). SgrS base pairs at a second site in the *manX-manY* intergenic region, 30 nucleotides upstream of the *manY* start codon. SgrS binding at this site represses *manY* and *manZ* translation (*manZ* translation is coupled to that of *manY*) by an unknown mechanism (Rice *et al.*, 2012). The *manY* binding site is outside the −20 to +20 (relative to the start codon) region protected by the translation initiation complex (Beyer *et al.*, 1994, Hüttenhofer & Noller, 1994, Bouvier *et al.*, 2008) suggesting that the mechanism of regulation by SgrS is not the canonical mechanism of occlusion of the SD sequence.

In this study, we probed the roles of Hfq, SgrS and the ribosome, particularly ribosomal protein (r-protein) S1, in modulating the translation of *manY*. We found that Hfq is required for SgrS-dependent regulation of *manY*, not only because it stabilizes SgrS (Balasubramanian & Vanderpool, 2013), but also because it promotes SgrS-*manY* mRNA duplex formation. We discovered that the AU-rich sequences that comprise the SgrS binding site upstream of *manY* act as a translational enhancer. The enhancer promotes higher levels of translation independent of whether the SD is strong or weak. We performed genetic and biochemical experiments that provide evidence for ribosomal protein S1 binding to the enhancer sequence to make the *manY* SD more accessible. Our data are consistent with the model that *manY* translation is controlled by an r-protein S1-dependent enhancer, and SgrS represses translation by interfering with S1 binding to the enhancer sequence, reducing translation efficiency.

## Results

### Hfq is essential for SgrS-mediated translational repression of *manY*

During glucose-phosphate stress, translation of the *manXYZ* operon is repressed by SgrS via base pairing interactions at two distinct sites on *manXYZ* mRNA (Fig. 1A) (Rice *et al.*, 2012, Rice & Vanderpool, 2011, Azam & Vanderpool, 2018). The base pairing interaction between SgrS and the *manX-manY* intergenic region (Fig. 1A, B) is 30 nucleotides upstream of the *manY* start codon (Rice *et al.*, 2012). Previous studies (Beyer *et al.*, 1994, Hüttenhofer & Noller, 1994, Bouvier *et al.*, 2008) have suggested that base pairing interactions outside the −20 to +20 window (relative to the start codon) cannot regulate translation by steric occlusion of ribosome binding, so the precise mechanism of SgrS-mediated translational repression of *manY* has remained unknown. To investigate the molecular details of the SgrS-*manY* interaction, we first looked at the role of the RNA chaperone Hfq (Møller *et al.*, 2002, Zhang *et al.*, 2002, Lease & Woodson, 2004). For these experiments, we used a *manY′-′lacZ* translational reporter fusion construct described previously (Rice & Vanderpool, 2011) that contains only the SgrS binding site in the *manX-manY* intergenic region (Fig. 1C). We monitored activity of the fusion in wild-type and Δ*hfq* host strains carrying SgrS expression plasmids. Previously, we showed that the stability of *E. coli* SgrS (SgrS*_Eco_*) is reduced in the absence of *hfq*, but *Salmonella* SgrS (SgrS*_Sal_*), which has a similar seed region and demonstrates comparable effectiveness in regulating targets in *E. coli*, is more stable in a Δ*hfq* background (Balasubramanian & Vanderpool, 2013). Consistent with our previous studies (Azam & Vanderpool, 2018, Balasubramanian & Vanderpool, 2013), in a wild-type host, both SgrS*_Eco_* and SgrS*_Sal_* repressed *manY′-′lacZ* fusion activity to less than 50 % of the activity in vector control cells (Fig. 1C). In the Δ*hfq* background, the basal level of *manY′-′lacZ* activity was reduced, and neither SgrS*_Eco_* nor SgrS*_Sal_* repressed translation of *manY′-′lacZ* further (Fig. 1C).

**Figure 1.**
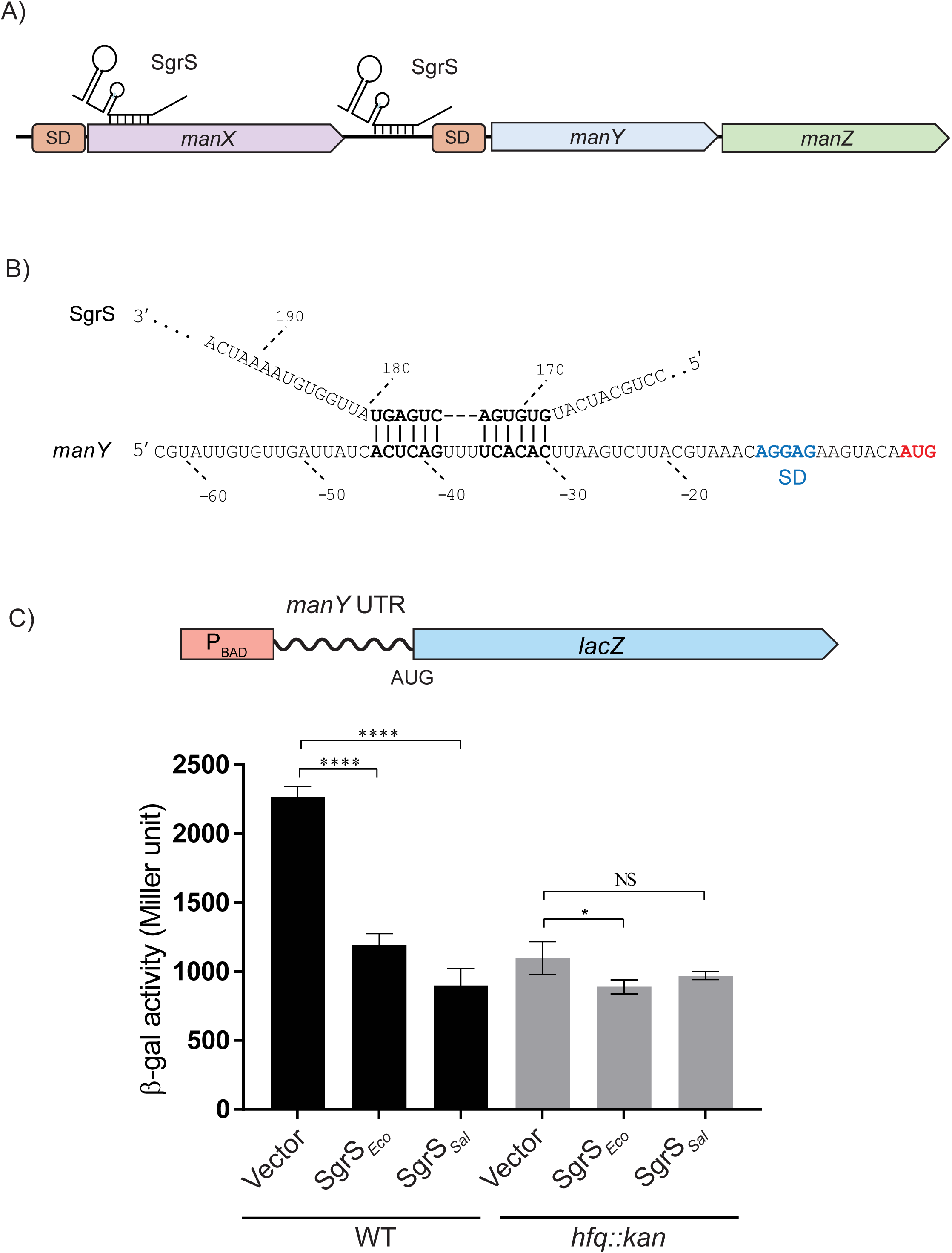
SgrS-dependent regulation of *manY* translation requires Hfq. A) A model showing the two SgrS binding sites on *manXYZ* mRNA. B) SgrS base pairing at the site in the *manX-manY* intergenic region, 30 nt upstream of the *manY* start codon. C) Wild-type (SA1747) and *hfq::kan* (SA1907) strains harboring a *manY′-′lacZ* translational fusion (under the control of a P_BAD_ promoter) were transformed with a vector or plasmid containing one of two SgrS orthologs – SgrS*_Eco_* from *E. coli*, or SgrS*_Sal_* from *Salmonella*. Cells were grown and β-Galactosidase assays were performed as described in the Materials and Methods. Error bars represent standard deviation from three biological replicates. *P* values (unpaired *t* test) are shown as: *, *P* < 0.05; **, *P* < 0.005; ***, *P* < 0.0005;****, *P* < 0.0001; NS, not significant.

To test whether Hfq is required to promote annealing between SgrS and *manY* mRNA, we performed footprinting reactions in the presence and absence of Hfq. End-labeled *manY* transcripts incubated with and without SgrS and Hfq were subjected to enzymatic (with RNase T1) and chemical (lead acetate) digestion. We performed an SgrS-*manY* footprinting experiment by mixing denatured end-labeled *manY* mRNA and unlabeled SgrS (with or without Hfq), followed by incubation under conditions that allow for annealing and then digestion. Using this procedure, we observed SgrS-dependent protection in the absence of Hfq (Fig. 2A, compare lanes 3 and 4) and in the presence of Hfq (Fig. 2A, lanes 5 and 6). The protection covered residues −30C to −44A (Fig. 2A, numbering relative to start codon), with hypersensitivity of the three U residues (−36U to −38U) in the center of the base pairing interaction that are not involved in duplex formation (Fig. 1B). These results are perfectly consistent with the SgrS binding site on *manY* mRNA that we previously mapped using genetic methods (Rice *et al.*, 2012).

**Figure 2.**
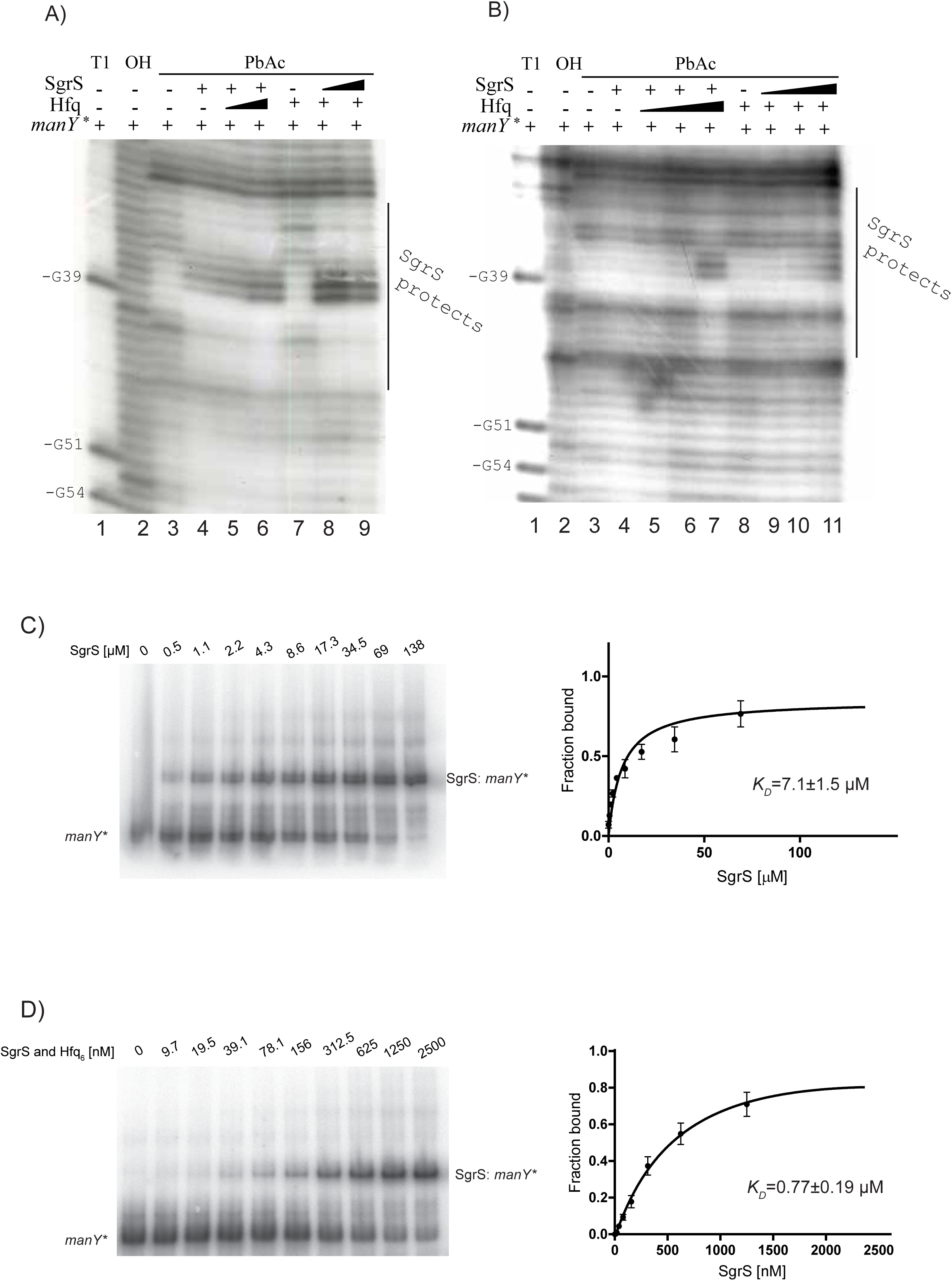
Hfq promotes duplex formation between SgrS and *manY* mRNA. A) Footprinting assays were performed using in vitro transcribed labeled *manY* mRNA mixed with SgrS and then denatured prior to addition of binding buffer and varying concentrations of Hfq. Lane 1 – “T1” – contains the G-ladder generated by RNase T1 digestion, Lane 2 – “OH” – contains the ladder generated by alkaline hydrolysis, and lanes 3-9 – “PbAc” – are lead acetate digested samples. Nucleotides protected by SgrS are labeled on the right. B) Footprinting assays were performed using in vitro transcribed – non-denatured – labeled *manY* mRNA mixed with SgrS and varying concentrations of Hfq. Lanes are labeled as described for part A. C) EMSAs using native gel electrophoresis were performed with P^32^ labeled *manY* and unlabeled SgrS that were hybridized in a binding buffer without prior denaturation. The samples were resolved in a chilled native acrylamide gel. Error bars represent standard deviation from (n=2) replicate experiments. D) EMSAs for SgrS and *manY* in the presence of Hfq. Hfq was removed by phenol extraction prior to running samples on a chilled native acrylamide gel. Error bars represent standard deviation from (n=3) replicate experiments. For both C and D, the measured band densities were plotted using the GraphPad Prism software to calculate the dissociation constants.

We next conducted footprinting experiments where SgrS and *manY* RNAs were not denatured prior to annealing and chemical probing (Fig. 2B). Under these conditions, we did not see SgrS-mediated protection of *manY* mRNA in the absence of Hfq (Fig. 2B, compare lanes 3 and 4) or when Hfq was present in the annealing mixture at lower concentrations (Fig. 2B lanes 5 and 6). In contrast, when Hfq was present at a higher concentration, we observed SgrS-dependent protection of *manY* mRNA over the known binding site – from residues −30C to −44A (Fig. 2B, lane 7). When reactions were performed using a limiting amount of Hfq and increasing amounts of SgrS (Fig. 2B, lanes 9, 10, and 11), we did not see protection. Thus, the requirement for Hfq to promote SgrS-*manY* mRNA interactions under these conditions could not be overcome by increasing concentrations of SgrS.

The data presented so far suggest that Hfq facilitates intermolecular SgrS-*manY* mRNA base pairing. To corroborate this finding, we conducted a set of electrophoretic mobility shift assays (EMSAs) to quantify SgrS-*manY* RNA interactions in the presence and absence of Hfq. For these experiments, in vitro transcribed *manY* and SgrS RNAs were mixed in a binding buffer without a prior denaturation step. We found that in the absence of Hfq, SgrS formed a complex with *manY* mRNA with a *K_D_* of 7.1 µM (Fig. 2C). When end-labeled *manY* RNA was incubated with increasing amounts of SgrS and equimolar concentrations of Hfq (which was subsequently removed by phenol extraction before the samples were resolved in native polyacrylamide gels), we obtained a *K_D_* of 0.7 µM, 10-fold lower than in the absence of Hfq (Fig. 2D). Altogether, the data suggest that Hfq promotes SgrS-mediated regulation of *manY* translation by facilitating SgrS-*manY* duplex formation.

### The *manY* translation initiation region contains a ribosome-dependent enhancer

SgrS acts as a translational repressor by binding at a site upstream of the two major determinants of ribosome binding – the Shine Dalgarno (SD) and initiation codon. We hypothesized that the region bound by SgrS might be a translation enhancer, similar to those described in mRNA targets of other sRNAs (Yang *et al.*, 2014, Sharma *et al.*, 2007). To test the role of the putative enhancer located within the SgrS binding site, we constructed a set of *lacZ* translational fusions. These fusions were driven by a heterologous promoter (P_BAD_), and contained the *manX-manY* intergenic region, including the SgrS binding site (Fig. 3A, green region). The wild-type fusion had a high basal level of activity at ∼4000 Miller Units (Fig. 3A, “WT”). The fusion lacking the SgrS binding site and putative enhancer had a greatly reduced basal level of activity (Fig. 3A, “Δenh”), consistent with the idea that this sequence has a stimulatory effect on *manY* translation. A third fusion where the putative enhancer was moved downstream of the ATG (maintaining the reading frame) had similarly low activity (Fig. 3A, “ATGenh”), suggesting that the translation stimulatory effect of this sequence is position-dependent.

**Figure 3.**
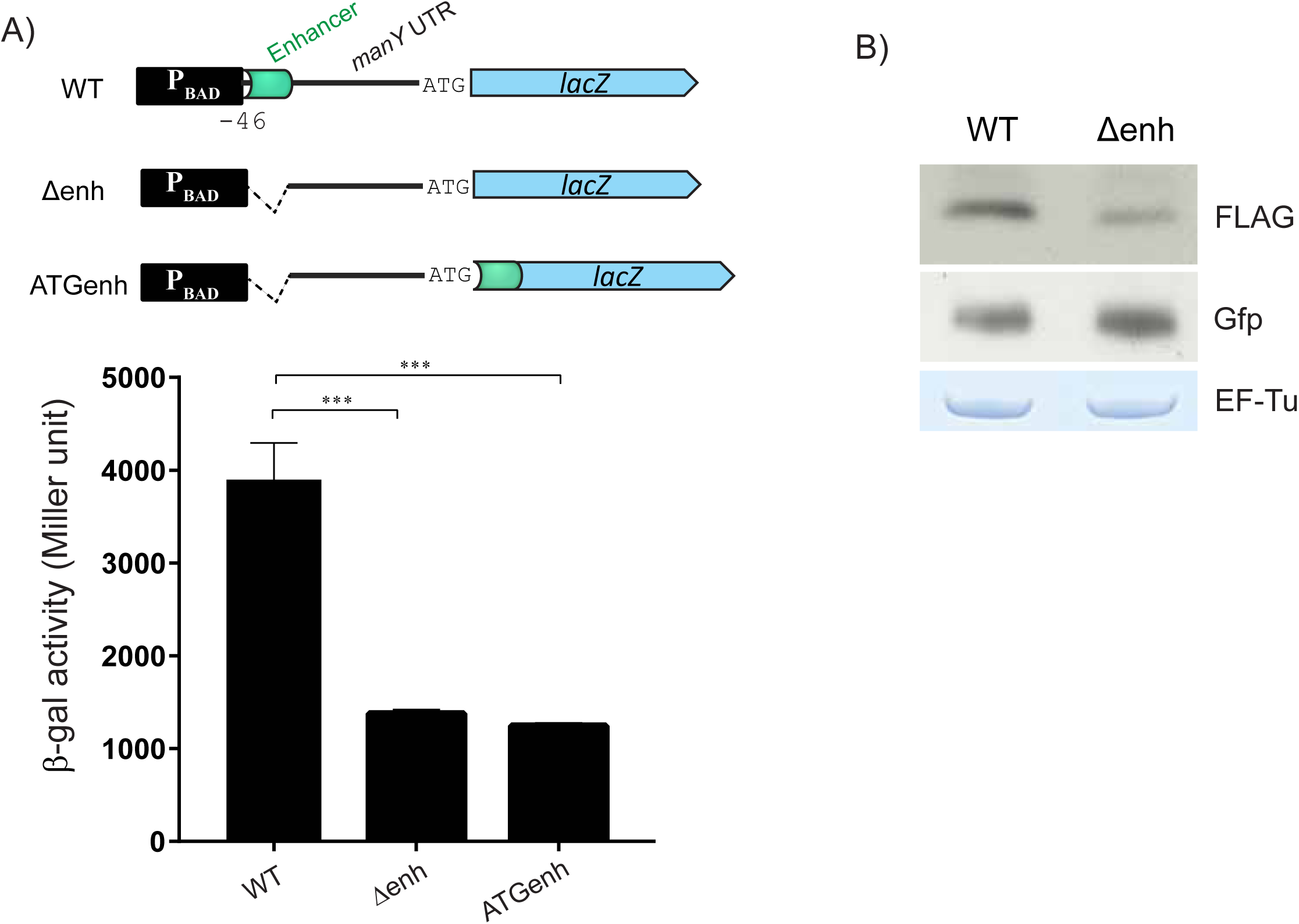
The translation enhancer is ribosome-dependent. A) Activities of three *manY′-′lacZ* fusions were compared to assess the regulatory role of the putative enhancer sequence. All fusions are controlled by the heterologous P_BAD_ promoter. “WT” contains the wild-type *manX-manY* intergenic region with the enhancer sequence (indicated by the green box) positioned ∼30 nt upstream of the *manY* start codon. “Δenh” contains a deletion of the enhancer sequence, “ATGenh” has the enhancer sequence deleted from the upstream region and replaced just downstream of (and in frame with) the start codon. Strains containing fusions were grown for 2 hr until reaching the mid-log phase and β-galactosidase assays were performed. Error bars represent standard deviation from three biological replicates. *P* values (unpaired *t* test) are shown as: *, *P* < 0.05; **, *P* < 0.005; ***, *P* < 0.0005;****, *P* < 0.0001; NS, not significant. B) Translation reactions were performed for 30 minutes using *manY-*3XFLAG transcripts. “WT” is wild-type enhancer sequence and “Δenh” contains a deletion of the enhancer sequence (as in part A). Products of translation were detected by Western blot using anti-FLAG and anti-GFP antibodies. As a control, reaction mixtures from each tube were analyzed on an SDS-polyacrylamide gel and stained with Coomassie Blue (the band for EF-Tu is shown).

To determine whether the enhancer activity of the region upstream of *manY* requires cellular factors other than the translation machinery itself, we performed *in vitro* translation assays with 3XFLAG-tagged *manY* transcripts. The amount of ManY-3XFLAG protein produced from wild-type transcripts containing the enhancer sequence in the 5’ UTR was substantially more than the protein produced from an equivalent amount of Δenh transcripts (Fig. 3B). These results suggest that the enhancer activity of the sequences upstream of *manY* is not dependent on Hfq, RNase E, or other cellular factors. Instead, we propose that the enhancer can stimulate translation of *manY* directly through interaction with the ribosome.

### Genetic analysis of enhancer sequences

The putative enhancer region is U-rich (Fig. 4A, −28 to −29 and −35 to −38), similar to the motifs found in previous studies that interact with the small ribosomal subunit (Duval *et al.*, 2013, Takahashi *et al.*, 2013, Marzi *et al.*, 2007, Boni *et al.*, 1991). We also noticed a “CACA” motif, similar to those reported to act as translation enhancers in other sRNA target mRNAs (Sharma *et al.*, 2007, Yang *et al.*, 2014). To further characterize the sequence determinants of the putative enhancer, we constructed reporter strains derived from the WT *manY′-′lacZ* fusion, each with different nucleotide substitution mutations in the putative enhancer region (Fig. 4A). Mutations in the U residues of the putative enhancer had the strongest effect on the basal levels of translation. Changing −37U and −38U to G residues (Fig. 4A, B, *mut1*) reduced activity by ∼30%. Mutation of −35U and −34U to G residues (Fig. 4A, B, *mut2*) had no effect on activity, but mutation of all four U residues to G residues (Fig. 4A, B, *mut3(G)*) reduced activity of the fusion by ∼85% compared to the wild-type fusion. Importantly, mutation of the U residues from −35 to −38 to A residues (Fig. 4A, B, *mut5(A)*) yielded a fusion with activity equivalent to the wild-type fusion. Mutation of U residues at positions −28 and −29 to G residues also had a strong impact on basal levels of activity (Fig. 4A, B, *mut4*). Other mutations in this putative enhancer region, including residues in the “CACA” motif, had no effect on basal levels of translation (Fig. 4A, B, *mut6, mut7, mut9, mut10, mut14, mut15*). These results suggest that U and A residues in this region are particularly important for the translation enhancing effect.

**Figure 4.**
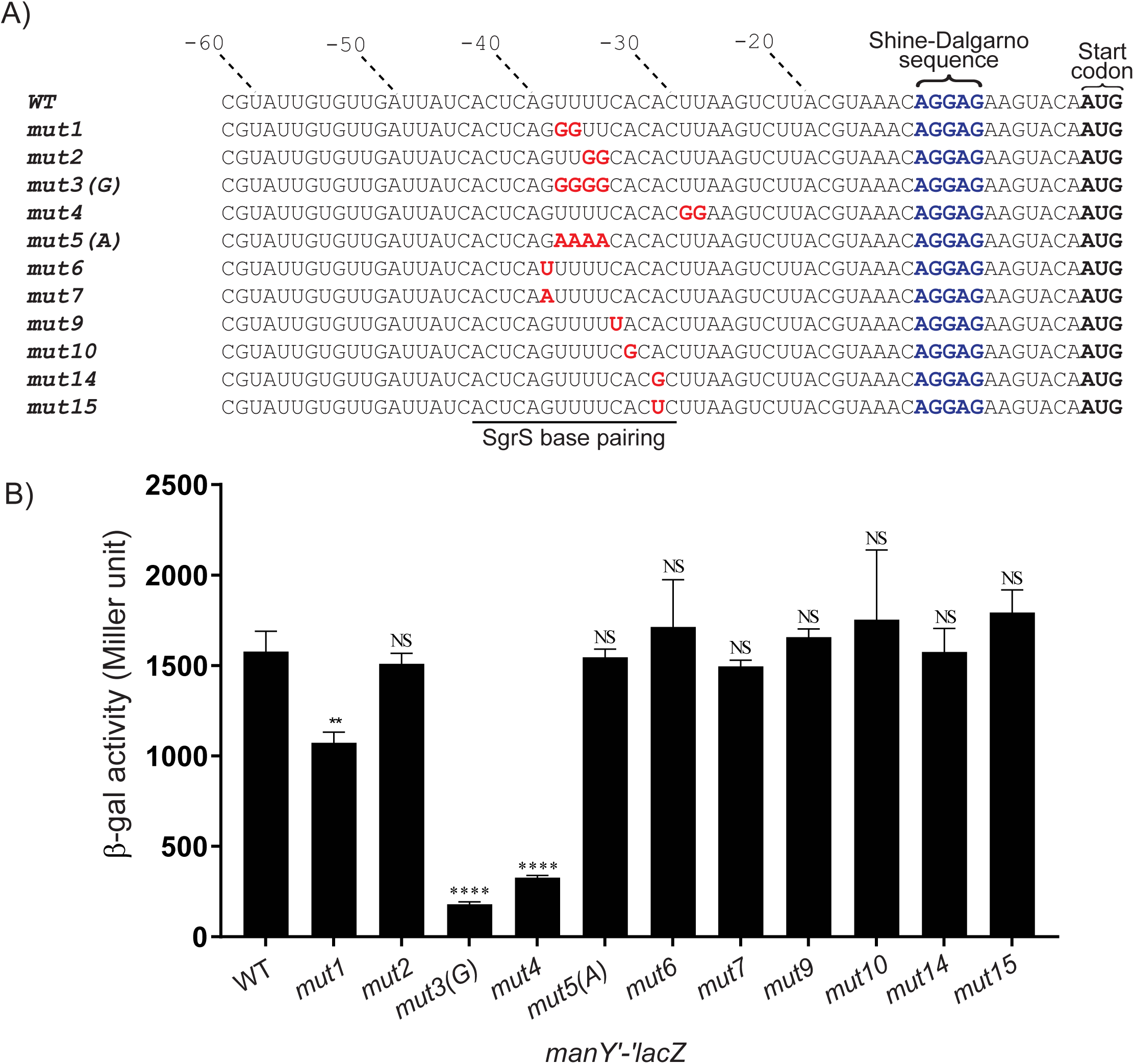
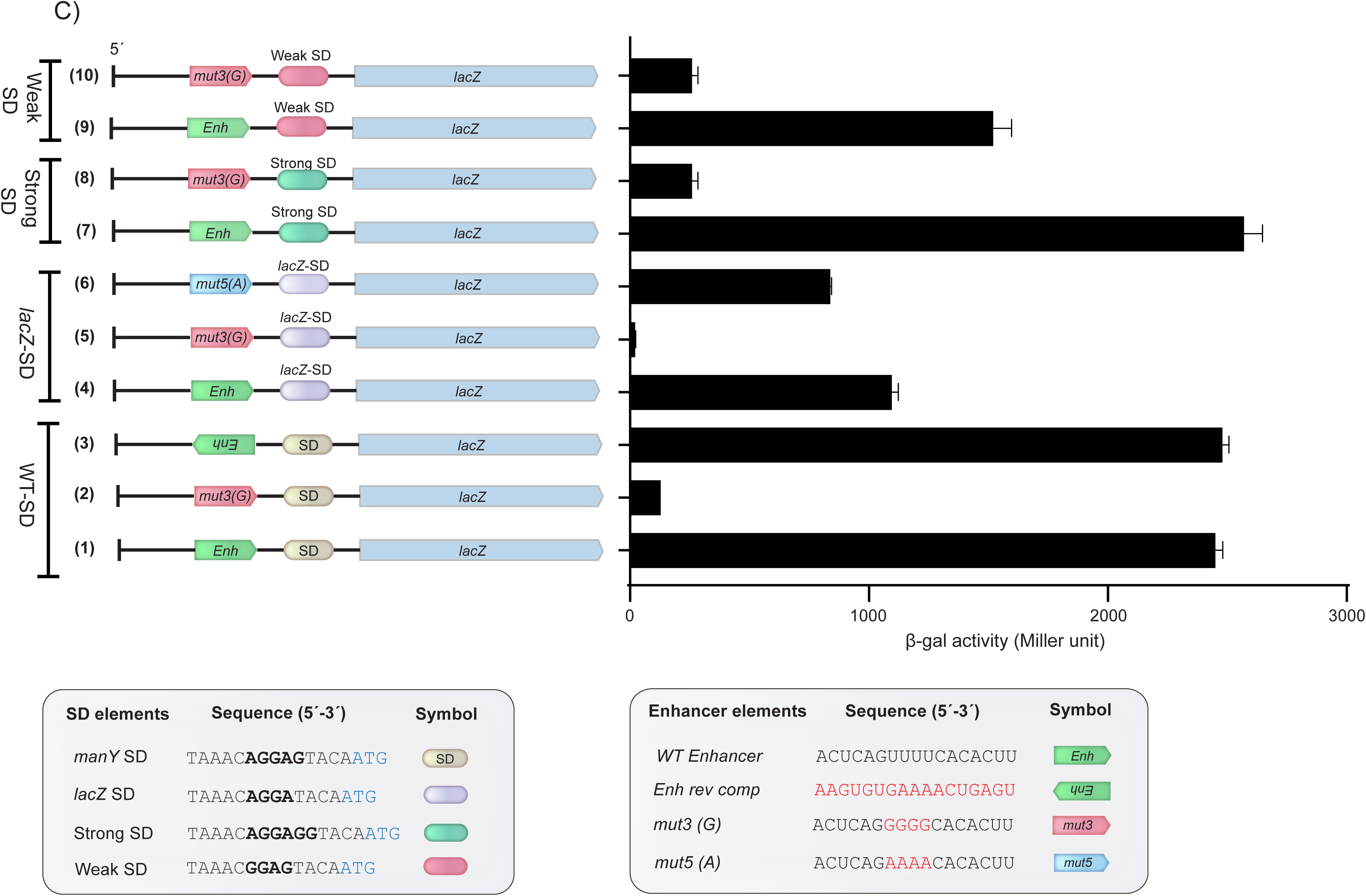
Mutational analyses of the AU-rich translation enhancer. A) A series of *manY* translational fusions containing wild-type or mutant enhancer sequences were assayed for β-Galactosidase activity. The top sequence (WT) is the wild-type sequence for the *manX-manY* intergenic region. The SD is indicated in blue and start codon in bold black letters. The sequences involved in base pairing with SgrS are indicated by the line under the sequences. Red letters in mutant sequences indicate positions of substitutions. B) β-Galactosidase assays of cultures containing indicated carrying the indicated fusions. Cells were grown for 2 hr until reaching the mid-log phase and β-galactosidase assays were performed. Error bars represent standard deviation from three biological replicates. *P* values (unpaired *t* test) are shown as: *, *P* < 0.05; **, *P* < 0.005; ***, *P* < 0.0005;****, *P* < 0.0001; NS, not significant. C) Indicated reporter fusions harbored in the degradosome mutant strain (*rne131::kan*) and assayed for β-Galactosidase activity as described in part B. D) *manY′-′lacZ* reporter fusions containing different combinations of SD and enhancer sequences were constructed and assayed for β-Galactosidase activity as described in part B. Symbols and corresponding sequences are defined in the insets below the graph. Fusions are labeled 1 through 10 and discussed in text accordingly. Error bars represent standard deviation from three biological replicates.

A number of studies have described AU-rich enhancer sequences located upstream of the start codon and SD, where r-protein S1 binds and enhances efficiency of translation initiation (Duval *et al.*, 2013, Takahashi *et al.*, 2013, Marzi *et al.*, 2007, Komarova *et al.*, 2005, Boni *et al.*, 1991). We hypothesized that the putative enhancer might similarly interact with S1 to promote translation of *manY*. If this were true, then the enhancer activity of this upstream sequence should be independent of SD-anti-SD interactions, and the enhancer should promote translation regardless of the strength of the SD. To test this possibility, we tested reporter constructs with various combinations of enhancer and SD sequences (Fig. 4C). As observed in the previous experiment, mutation of U residues to G residues (*mut3(G)*) strongly reduced the activity of the reporter fusion with the native *manY* SD (Fig. 4C, compare activity of reporters 1 and 2). A construct containing the native *manY* SD and the reverse complement of the enhancer region – now A-rich rather than U-rich – had high levels of translation (Fig. 4C, reporter 3), comparable to the wild-type reporter. This result suggests that the translation-enhancing effect of this region is not strictly sequence-dependent, but that AU-rich sequences in general are stimulatory. This is consistent with the known sequence preferences of r-protein S1.

We next tested whether the enhancer could stimulate translation from a weaker SD. The native *manY* SD (5’-AGGAG-3’) was swapped for the slightly weaker *lacZ* SD (5’-AGGA-3’), leaving the rest of the sequence in the reporter the same. As expected, the activity of this reporter was reduced compared to the reporter with the stronger *manY* SD (Fig. 4C, compare activity of reporter 1 and reporter 4). The *mut3(G)* enhancer sequence strongly reduced activity (Fig. 4C, reporter 5), while the *mut5(A)* enhancer sequence restored the activity (Fig. 4C, reporter 6) of the constructs with the *lacZ* SD. The same patterns were seen for two additional SD variants (Fig. 4C, reporters 7, 8, 9 and 10). Regardless of the SD sequence, the wild-type U-rich enhancer promoted higher levels of translation, and mutation of U residues to Gs dramatically reduced the translation activity. These results suggest that the enhancer functions independent of the SD to promote translation.

### Ribosome protection of the non-contiguous RBS and translation enhancer

The analyses presented so far suggest that an upstream enhancer sequence plays a positive role in *manY* translation. We hypothesized the enhancer interacts with the initiating ribosome, specifically r-protein S1, and that this interaction increases translation initiation. Studies of *thrS* mRNA, encoding threonyl-tRNA synthetase, showed that the 5’ UTR of *thrS* contains a bipartite ribosome binding site where an upstream AU-rich element interacts with r-protein S1 ((Duval *et al.*, 2013, Romby *et al.*, 1996), Fig. 5A). The AU-rich S1 binding site acts as a translational enhancer for *thrS* mRNA (Duval *et al.*, 2013, Takahashi *et al.*, 2013, Marzi *et al.*, 2007, Boni *et al.*, 1991). We used this mRNA as a comparison and positive control in the next experiments to further examine the role of S1 in regulation of *manY* translation. We made *thrS* translational fusions with the wild-type sequence and mutants where U residues were changed to either Gs or As (Fig. 5A). β-galactosidase assays showed the same pattern we observed for the *manY* fusions. The wild-type *thrS* fusion (Fig. 5B, “WT-*thrS*”) had a high level of activity, which was strongly reduced in the mutant fusion with G residues (Fig. 5B, “G-*thrS*”) and restored in the mutant fusion with A residues (Fig. 5B, “A-*thrS*”). Footprinting reactions performed using RNase T1 (enzymatic) and lead acetate (chemical) probing with end-labeled mRNA and purified *E. coli* ribosomes showed the expected bipartite RBS for *thrS* mRNA, with protection around the SD and at the upstream AU-rich enhancer (Fig. 5C). We then performed footprinting reactions with end-labeled *manY* mRNA and purified ribosomes. In the absence of ribosomes, the pattern of cleavage by lead acetate suggested that the sequences around the *manY* SD and start codon are not very accessible and may be sequestered in a secondary structure. This is consistent with other examples of S1-dependent translation of mRNAs with highly structured translation initiation regions (Boni *et al.*, 2001, Romilly *et al.*, 2019). In the presence of ribosomes, we observed a bipartite footprint, similar to the *thrS* footprint, with protection of the SD and start codon as well as protection of the upstream region containing the U-rich enhancer (Fig. 5D). These results are consistent with our hypothesis that the *manY* leader contains an enhancer upstream of the SD, which interacts with the ribosome, possibly with r-protein S1.

**Figure 5.**
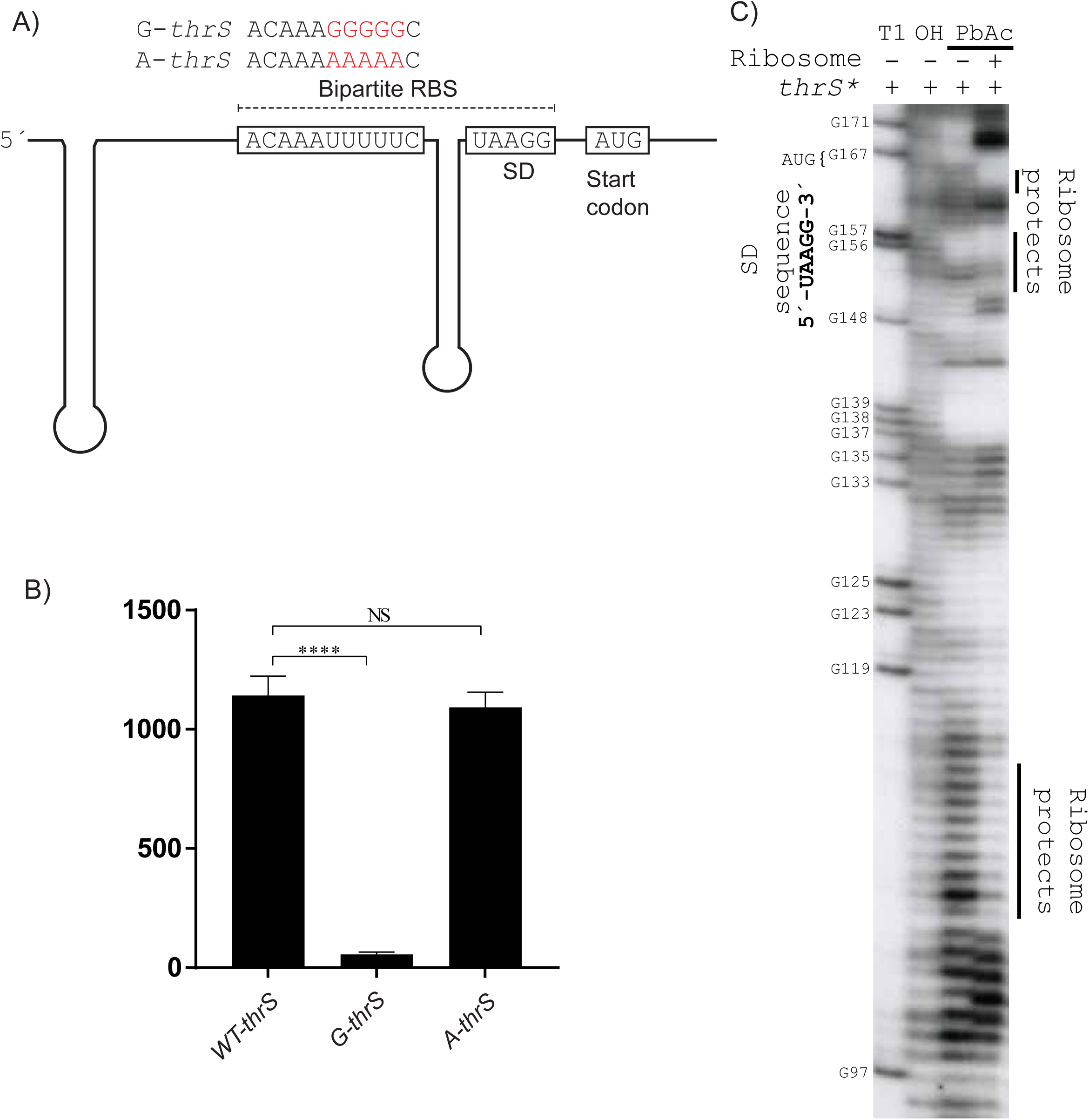
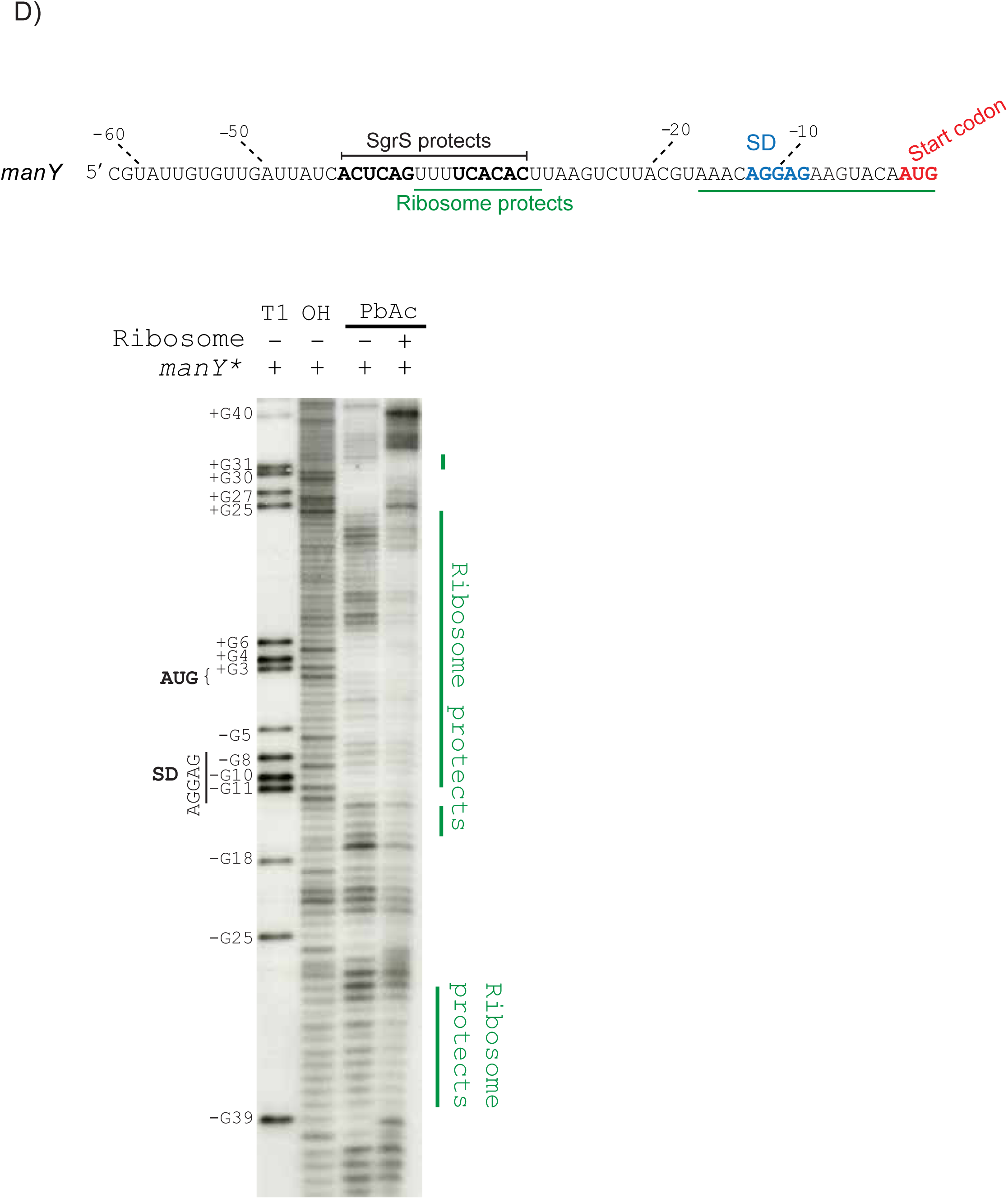
Like known r-protein S1 target *thrS*, the *manY* leader contains a bipartite RBS. A) Graphical representation of *thrS* mRNA, including the bipartite RBS. The AU-rich region upstream of the SD is known to encompass a binding site for r-protein S1. Substitution mutations in the S1 binding site are indicated above in red nucleotides – “*G-thrS*” and “*A-thrS*.” B) *thrS′-′lacZ* translational fusions were constructed. “WT-*thrS*” indicates the wild-type fusion, “*G-thrS*” and “*A-thrS*” are the mutant fusions with sequences as indicated in part A. Error bars represent standard deviation from three biological replicates. *P* values (unpaired *t* test) are shown as: *, *P* < 0.05; **, *P* < 0.005; ***, *P* < 0.0005;****, *P* < 0.0001; NS, not significant. C) Ribosome footprint analysis of *thrS* mRNA. In vitro transcribed mRNA was end-labeled with ^32^P and incubated with and without *E. coli* ribosomes. Lanes are labeled as described in Fig. 2A legend. Positions of the start codon and SD are indicated on the left. Ribosome protected nucleotides are indicated on the right. D) The *manY* UTR and early coding region sequences are shown. The sequences comprising the SgrS binding site and enhancer are indicated with a line above the sequence. The SD and start codon are indicated in blue and red print, respectively. The regions protected by the ribosome are indicated with a green line below the sequences. The ribosome footprint is labeled as in part C.

### Ribosomal protein S1 binds to the *manY* 5’ UTR

A large body of literature has characterized S1 RNA-binding activity (Yokota *et al.*, 1979, Laughrea & Moore, 1977, Suryanarayana & Subramanian, 1979, Subramanian, 1983). In gram-negative bacteria, S1 orthologs are composed of six S1 motifs that are structurally similar (Fig. 6A), but diverge in amino acid sequence (Subramanian, 1983, Salah *et al.*, 2009). A cryo-EM based approach found that the first two motifs of S1 at the N-terminus interact with the r-protein S2 and function as a platform for ribosome binding (Loveland & Korostelev, 2018). Using proteins with mutations in the S1 motifs, the Marzi group determined that the first three N-terminal domains are crucial for S1 binding to *rpsO* mRNA (Duval *et al.*, 2013). Several residues of the third motif (Y205, F208, H219, and R254) were found to be crucial for interaction with Qβ RNA (Takeshita *et al.*, 2014). We aligned S1 proteins from 11 organisms belonging to the Proteobacteria and found that three out of these four residues important for RNA binding (Y205, F208, H219) were highly conserved (Fig. 6B). Alignment of S1 motifs from seven other diverse *E. coli* RNA binding proteins revealed that residues F208 and H219 were still fairly well conserved (Fig. 6C). These results led us to test whether mutation of conserved residues would impact S1 RNA binding activity. To address this question, we performed EMSAs with purified wild-type S1 and mutant S1 (mS1, Y205A, F208A, H219A) proteins and in vitro transcribed *thrS* and *manY* RNAs. Wild-type S1 bound to *thrS* mRNA with a *K_D_* of 3.1 nM, while mS1 had a higher *K_D_* of 16 nM (Fig. 6D). This result is consistent with the idea that mS1 is slightly defective for binding a known target. The interaction between wild-type S1 and *manY* mRNA was weaker than for *thrS*, with a *K_D_* of 81 nM. As for *thrS*, the interaction between mS1 and *manY* RNA was diminished (*K_D_* = 590 nM, Fig. 6E). Though higher than for *thrS* mRNA, these dissociation constants for *manY* mRNA are in a similar range to those reported previously for other S1 targets (Qureshi *et al.*, 2018). Altogether, the data suggest that the third S1 motif is important for binding of *thrS* and *manY* mRNAs.

**Figure 6.**
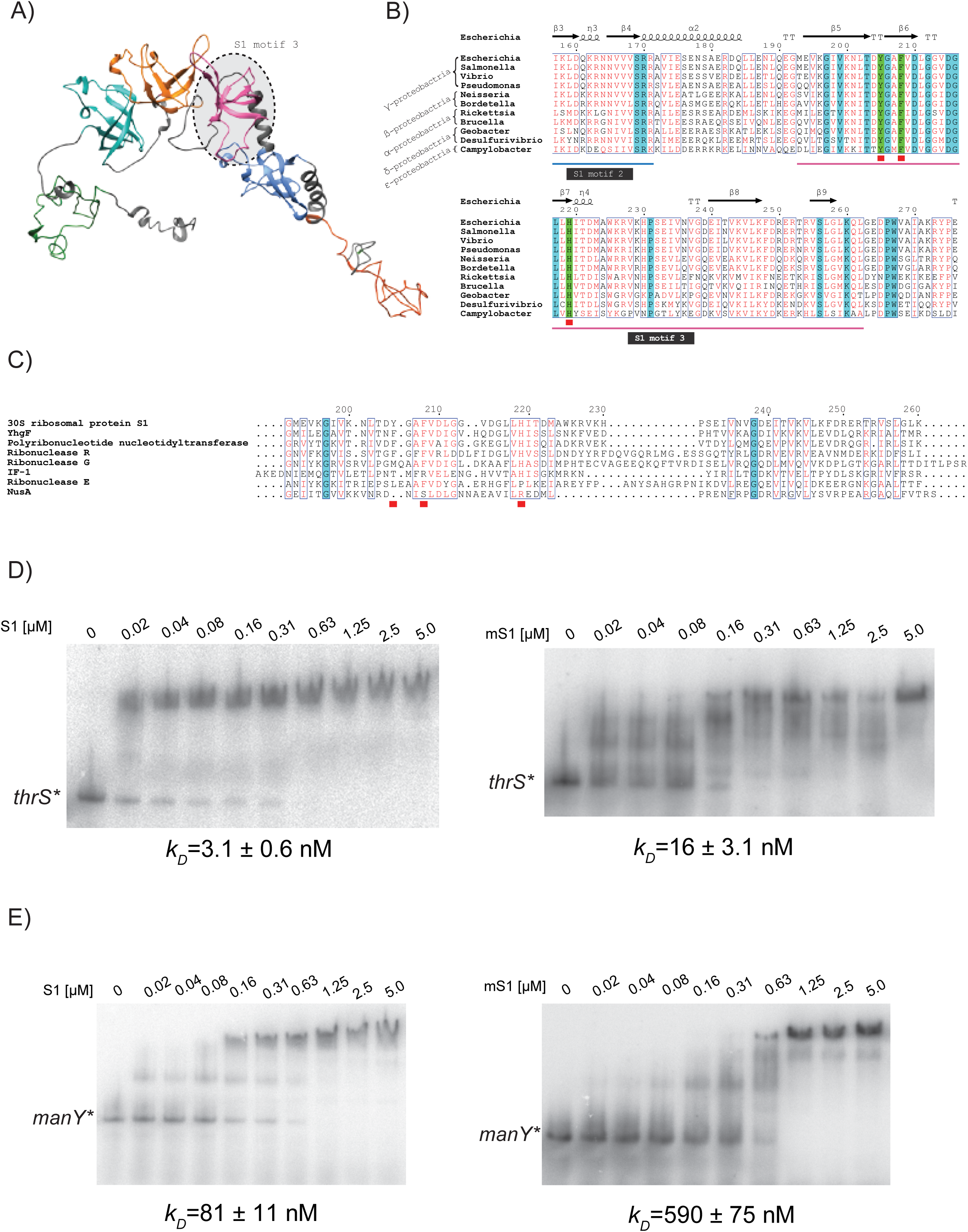
Characterization of wild-type and mutant S1 (mS1) protein binding to *thrS* and *manY* mRNAs. A) A structure of S1 (showing six S1 motifs, the N-terminal motif is on the right) predicted using I-TASSER homology modeling algorithm based on the deposited cryo-EM structure (Beckert *et al.*, 2018). S1 motif 3 is indicated with a dotted circle. B) A global alignment of S1 motif 3 from diverse bacterial species. The alignment was calculated using the Clustal Omega algorithm (Madeira *et al.*, 2019) and represented using the ESPript 3 web tool (Robert & Gouet, 2014). α-helices are displayed as squiggles. β-strands are marked with arrows, strict β-turns with the letters TT. The η symbol was used for 3_10_-helices. C) A global alignment of S1 motif 3 with amino acid sequences of S1 motifs from other *E. coli* RNA binding proteins. D-E) In vitro transcribed *thrS* and *manY* mRNAs were labeled at the 5’-end with P^32^ and mixed with WT S1 or mS1 in a binding buffer for EMSAs. Band intensities were measured for three replicates to determine the dissociation constants.

To further study the interaction of r-protein S1 and *manY* mRNA, we performed additional footprinting reactions using diethyl pyrocarbonate (DEPC), which primarily modifies exposed adenine residues (Ehresmann *et al.*, 1987). After treatment with DEPC, a reverse transcription reaction using a labeled primer is performed. Bands represent positions that were DEPC-modified, causing reverse transcriptase to stop. In the absence of S1 protein, there was very little modification of A residues around the SD and start codon (Fig. 7, lane 2), again suggesting that the *manY* mRNA translation initiation region might be sequestered in a secondary structure. In the presence of wild-type S1 protein, there was substantially more modification, particularly in the region between the enhancer and the SD. This suggests that S1 binding to this region might remodel the secondary structure to make it more accessible for formation of the translation initiation complex. In the presence of mS1, we also saw new modifications signifying structural rearrangements, but these were distinct from the changes seen in the presence of wild-type S1 (Fig. 7). This result is consistent with the EMSA (Fig. 6E) and suggests that mS1 binds differently (and more weakly) to the *manY* mRNA.

**Figure 7.**
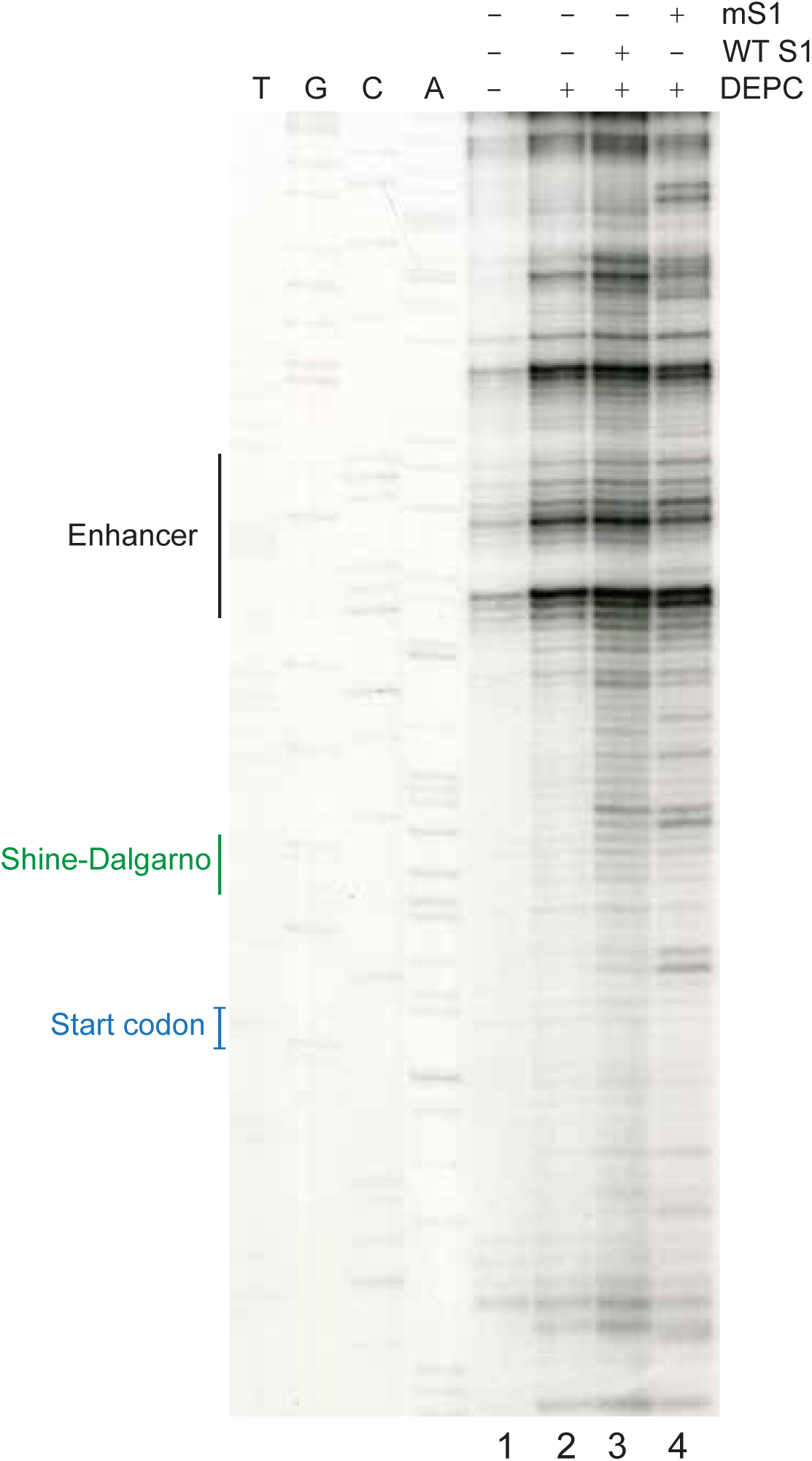
S1 unfolds the secondary structures in the *manY* UTR. The *manY* transcript was preincubated with S1 or mS1 before chemical modification. DEPC modified *in vitro* transcribed *manY* transcripts were used as a template for primer extension with a reverse transcriptase in the absence or presence of the protein (S1 and mS1), and resolved in a polyacrylamide-urea gel. Sanger ladders (T, G, C, and A) were generated with dideoxy-sequencing. The enhancer region, the SD, and the start codon are indicated on the left.

### Evidence of SgrS competition with S1 for control of *manY* translation

We hypothesized that SgrS represses *manY* translation by competing with r-protein S1 for binding at the enhancer sequence. To test this hypothesis in vivo, we constructed plasmids for ectopic expression of S1 and mS1, reasoning that a brief pulse of expression could alter the competition between endogenous SgrS and S1 protein. Previous work demonstrated that S1 stimulates translation of target genes in a narrow range of concentrations – at higher concentrations, S1 represses target translation (Boni *et al.*, 2001, Delvillani *et al.*, 2011). Since *manY* and *thrS* have different affinities for S1, we reasoned that S1-dependent stimulation of translation would require different levels of induction in vivo. Thus, we made two different types of S1 expression constructs – one with a strong SD sequence, and one with a weak SD sequence. We transformed plasmids with S1 or mS1 into strains harboring *thrS′-′lacZ* (positive control), *sodB′-′lacZ* (negative control), and *manY′-′lacZ*, and performed β-galactosidase assays after a brief induction. Since *thrS* has a higher affinity for S1, we used expression constructs with the weak SD. In the *thrS′-′lacZ* strain, ectopic production of wild-type S1 resulted in a very slight, but not statistically significant increase in β-galactosidase activity. Production of mS1 had a more pronounced effect – decreasing the activity by 16% (Fig. 8A). This result is consistent with the idea that the ectopically produced mS1 protein interferes with endogenous S1 activity and inhibits S1-mediated translational activation of a known target. In the strain with negative control *sodB′-′lacZ*, we used S1 and mS1 constructs with the strong SD sequence. There was no difference in β-galactosidase activity between control and S1- or mS1-producing strains (Fig. 8B).

**Figure 8.**
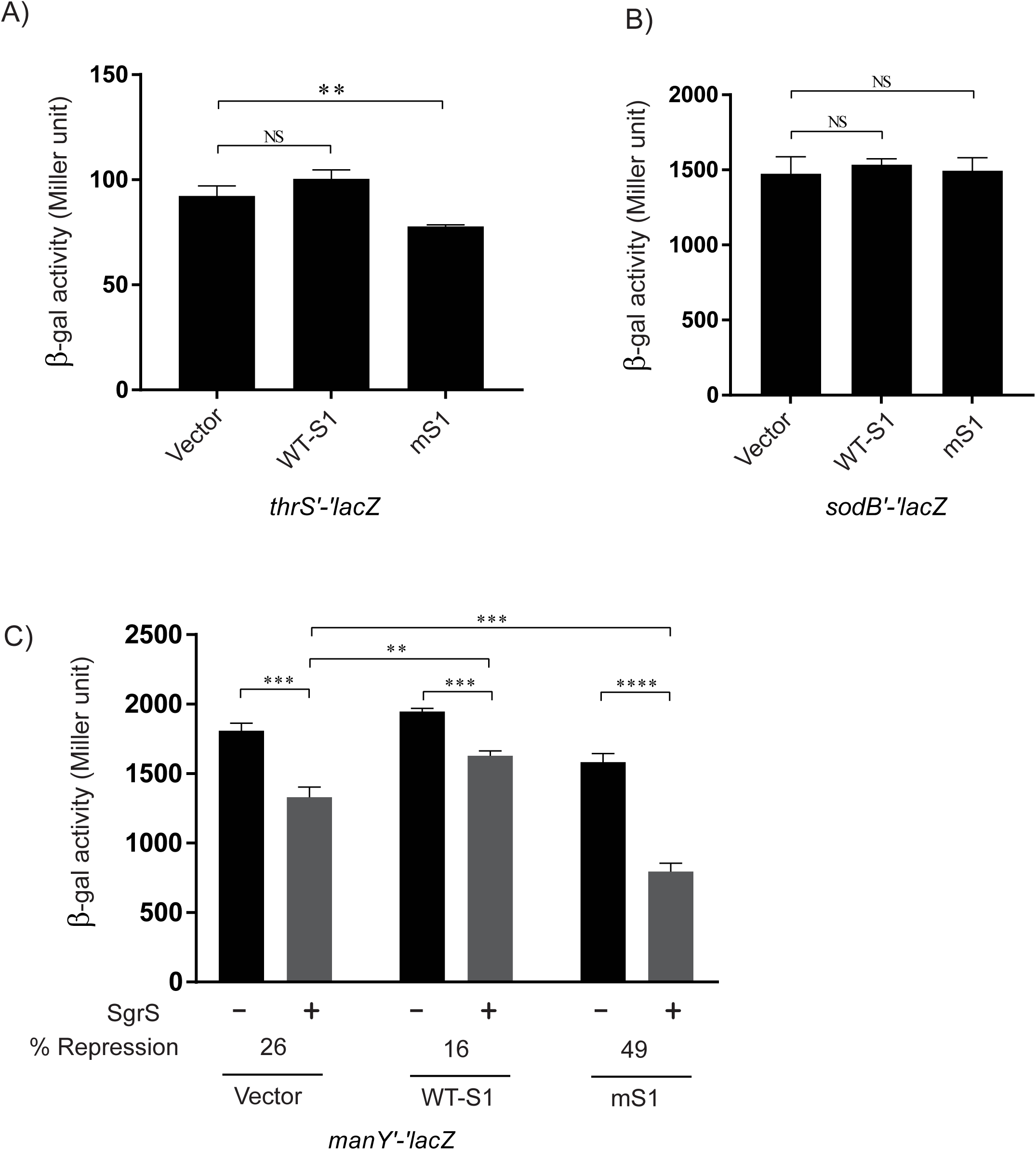
Short-term ectopic production of S1 or mS1 modulates the efficiency of SgrS-dependent translational regulation of *manY.* Low copy plasmids carrying *rpsA* encoding wild-type or mutant r-protein S1 under the control of an IPTG-inducible promoter were transformed into strains with *thrS′-′lacZ* (A), *sodB′-′lacZ* (B) or *manY′-′lacZ* (C) fusions (reporter fusions were under the control of arabinose-inducible P_BAD_ promoter). Cultures were grown at 37°C to OD_600_∼0.4 and the reporter fusion was induced with L-arabinose. Cells were grown for another 20 minutes before IPTG (to induce S1 or mS1) was added. Cultures were grown for an additional 10 min before β-galactosidase assays were performed. For *manY′-′lacZ* (C) fusions, SgrS and S1 (or mS1) was induced instantaneously. Error bars represent standard deviation from three biological replicates. *P* values (unpaired *t* test) are shown as: *, *P* < 0.05; **, *P* < 0.005; ***, *P* < 0.0005;****, *P* < 0.0001; NS, not significant.

In the *manY′-′lacZ* strain, we used the S1 and mS1 constructs with the strong SD sequence. We also performed the experiments in the absence and presence of α-methyl glucoside, to induce production of SgrS (Vanderpool & Gottesman, 2004). In the absence of SgrS, we saw the same S1-dependent patterns of activity as for *thrS*. Production of wild-type S1 slightly increased the activity of *manY′-′lacZ* and production of mS1 had an inhibitory effect (Fig. 8C, compare the black bars). As predicted if SgrS and r-protein S1 compete for binding to *manY* mRNA, the efficiency of SgrS-dependent repression varied depending on S1 production. In the strain with the vector control, SgrS induction caused a 26% reduction in β-galactosidase activity (Fig. 8C). (Note, this is a more modest repression ratio than seen in other experiments because we used a much shorter time for SgrS induction.) In the strain producing wild-type S1, the SgrS-dependent reduction in activity was only 16%. This suggests that wild-type S1 overproduction can modestly protect *manY* from SgrS-dependent translational repression. Most strikingly, production of mS1 left *manY* more susceptible to SgrS-dependent repression, with a repression value of 49% (Fig. 8C). These data are all consistent with the model that r-protein S1 stimulates *manY* translation, and that SgrS binding inhibits translation by interfering with S1-dependent activation.

## Discussion

In this report, we show that the *E. coli* sRNA SgrS, in collaboration with the RNA chaperone Hfq, represses *manY* translation by base-pairing with sequences comprising a translational enhancer. The enhancer stimulates *manY* translation in vitro, suggesting that the effect is mediated by ribosomes and not by other cellular factors. The AU-rich enhancer increases translation regardless of the strength of the SD sequence. The elements within the *manY* leader region resemble other so-called bipartite RBSs where ribosomes interact with both the SD and start codon region as well as a non-contiguous upstream region. Our data implicate r-protein S1 as an important mediator of *manY* translation initiation via interactions with the AU-rich enhancer sequence. Our data are consistent with a model where SgrS competes with S1 for binding to the enhancer sequences, thereby reducing S1-dependent translation and resulting in the observed SgrS-dependent repression of *manY* (Fig. 9).

**Figure 9.**
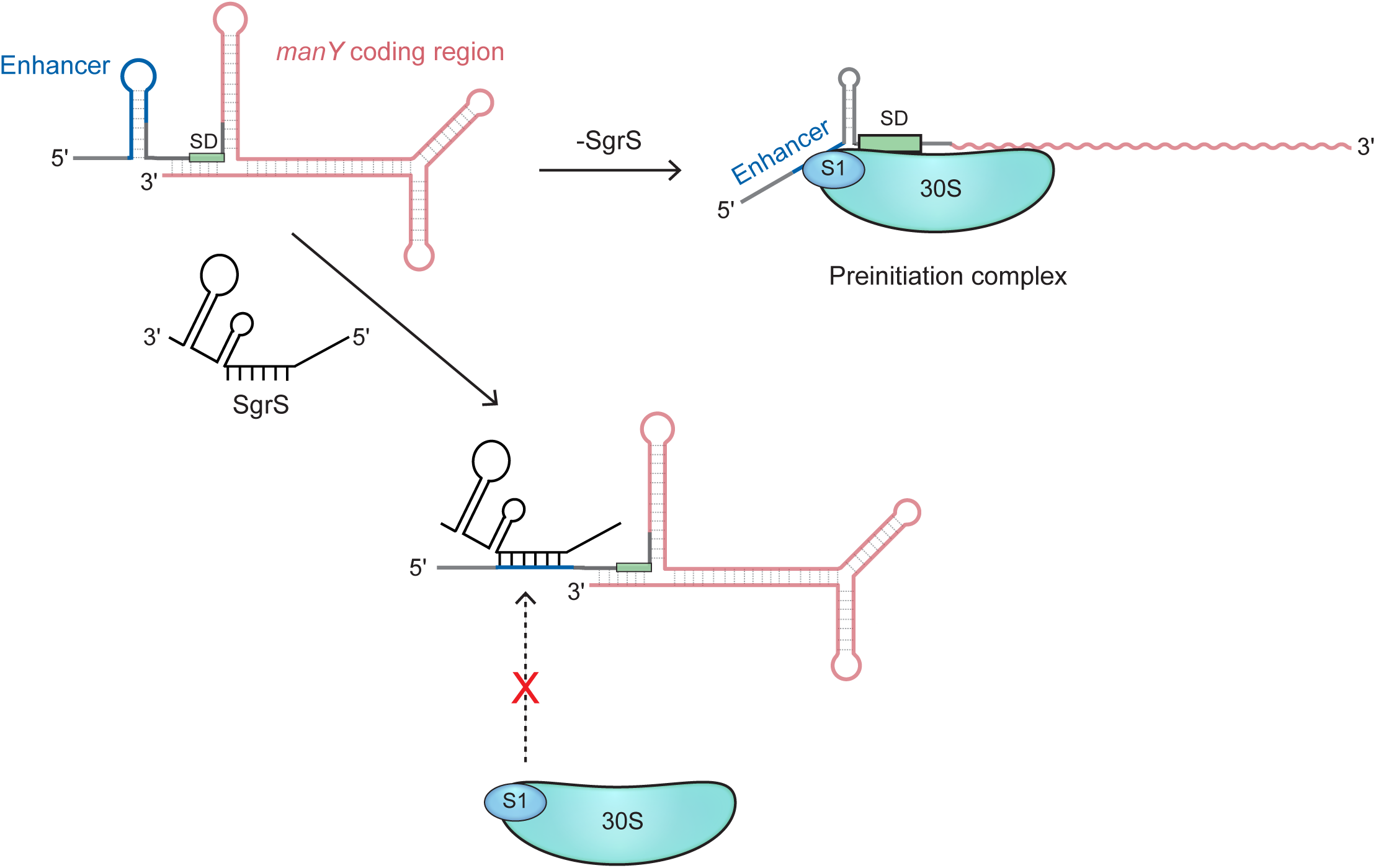
Model of SgrS-mediated enhancer silencing. Our results suggest that the *manY* translation initiation region contains structural elements that are remodeled in part by r-protein S1 binding to a translational enhancer upstream of the *manY* SD to promote high levels of translation in the absence of the sRNA SgrS. Under glucose-phosphate stress conditions, when SgrS is produced, it base pairs with sequences in the enhancer, preventing formation of the translation initiation complex. SgrS could inhibit translation by sequestering the enhancer and preventing binding of r-protein S1 or by altering a crucial the secondary structure that interacts with ribosome-bound S1.

The RNA chaperone Hfq is essential for *manY* translation repression by SgrS in vivo (Fig. 1C). Hfq is a pleiotropic global regulator of gene expression, owing to its dynamic and complex interaction patterns with many cellular RNAs (Vogel & Luisi, 2011, Woodson *et al.*, 2018, Melamed *et al.*, 2016). The involvement of Hfq in sRNA-mediated translational repression has been attributed to its ability to remodel RNA secondary structures and overcome energetic barriers presented by one or both RNA partners, and to enhance local concentrations of potential binding partners to increase the rates of association (Santiago-Frangos & Woodson, 2018). A recent report illustrated an example of Hfq-dependent structural remodeling of *dgcM* mRNA to allow duplex formation with OmrA or OmrB sRNAs, which results in translational repression of *dcgM* (Hoekzema *et al.*, 2019). Similar mechanisms have been found for other sRNA-mRNA pairs that require Hfq-dependent structure remodeling (Wroblewska & Olejniczak, 2016, Chen & Gottesman, 2017). In this study, Hfq was required in vivo for SgrS-dependent regulation of *manY* (Fig. 1C). In vitro, the presence of Hfq promoted SgrS-dependent protection of *manY* mRNA (Fig. 2A, B), and adding Hfq to in vitro annealing reactions resulted in a 10-fold reduction in SgrS-*manY K_D_* (Fig. 2C, D). However, we did not observe Hfq-mediated structural changes in our footprint experiments (Fig. 2A, B). We hypothesize that SgrS sequesters the enhancer from ribosome-associated S1 protein (Fig. 9). This model could explain why Hfq is essential for SgrS-mediated silencing of the enhancer in vivo. S1 forms a complex with *manY* with a *K_D_* of 80.5 nM (Fig. 6E) while SgrS forms a complex at a higher stoichiometry (*K_D_* of 7.1 µM, Fig. 2C). In the presence of Hfq, the SgrS-*manY* complex has a submicromolar dissociation constant (*K_D_* of 0.77 µM, Fig 2E), which could allow SgrS to effectively compete with S1 for binding to the same site on *manY* mRNA.

The *manY* mRNA leader possesses a bipartite RBS. The RBS of a gene often extends beyond the start codon and the SD (Shine & Dalgarno, 1974, Kozak, 1999, Steitz & Jakes, 1975) with a non-contiguous architecture where additional sequences in the 5’ leader establish interactions with ribosomal proteins and modulate translation initiation complex formation (Kudla *et al.*, 2009, Del Campo *et al.*, 2015, Nivinskas *et al.*, 1999, Tuerk *et al.*, 1988, Gold, 1988). The threonyl-tRNA synthetase (*thrS*) mRNA contains a bipartite RBS, with an upstream single-stranded U-rich enhancer that is separated from the SD region by a stem-loop structure. For *thrS* mRNA, the enhancer is recognized by r-protein S1 (Sacerdot *et al.*, 1998). This is just one example of an S1 binding site, and there are many other varieties (Sacerdot *et al.*, 1998, Duval *et al.*, 2013, Boni *et al.*, 2001, Romilly *et al.*, 2019), making it difficult to identify S1 binding sites based on sequence or structural characteristics. The *manY* enhancer is centered at ∼25 nt upstream of the SD and we see a bipartite pattern of ribosome-dependent protection (Fig. 5D) similar to the pattern for known S1 target *thrS* mRNA (Fig. 5C). In the absence of ribosomes, the *manY* SD and start codon region is not accessible to cleavage by lead acetate (Fig. 5D), suggesting it is structured. This is common for mRNAs whose translation is activated by r-protein S1, as S1 binding upstream can promote unfolding of secondary structure to allow formation of ribosome preinitiation complexes (Qureshi *et al.*, 2018, Kolb *et al.*, 1977, Bear *et al.*, 1976). Our DEPC modification and footprinting experiments also showed that the SD and start codon region of *manY* mRNA is rather inaccessible to modification, and that addition of S1 slightly enhances accessibility of these sequences (Fig. 7). Consistent with our results, other studies have demonstrated S1-mediated unfolding of secondary structures with free S1 proteins (Qureshi *et al.*, 2018, Bear *et al.*, 1976). In an elegant study, the Marzi group demonstrated that ribosome-bound S1 could unfold *rpsO* mRNA and expose the SD region more efficiently than free S1 protein (Duval *et al.*, 2013). Another recent study showed how ribosome-associated S1 prevents secondary structure formation and exposes an otherwise inaccessible SD of the *tisB* mRNA (Romilly *et al.*, 2019). With assistance from neighboring ribosomal components, S1 might play a more prominent role in remodeling the secondary structure of *manY* mRNA in its natural ribosome context. Nevertheless, our data are consistent with a model where *manY* translation is enhanced by r-protein S1 binding to a region upstream of the SD in the absence of SgrS (Fig. 9).

Both technical challenges and opportunities continue to arise as we discover more and more examples of sRNA-mediated translational regulation from a distance, *i.e.*, outside the boundaries where canonical steric occlusion of the SD and start codon are possible. There are many mRNAs with very long and highly structured 5’ leaders, and doubtless an array of new regulatory mechanisms controlling mRNA translation and stability are yet to be discovered. Our study and the numerous recent global analyses of RNA-RNA interactomes (Desnoyers & Massé, 2012, Azam & Vanderpool, 2018, Sharma *et al.*, 2007) suggest that there are dozens of targets for most sRNAs, and we must expand the search window beyond the SD and start codon region when looking for sRNA target binding sites. Mechanistic characterization of 5’ UTR-targeting sRNAs could reveal new functional elements involved in translation initiation and help us uncover the complex and dynamic interactions between mRNAs and ribosomes.

## Experimental Procedures

### Strains and plasmids

The strains, plasmids, and oligonucleotides used in this study are listed in Tables S1 and S2. *E. coli* K12 MG1655 derivatives were used for all experiments. P1 transduction (Miller, 1972) or λ-red recombination (Yu *et al.*, 2000) procedures were used to move alleles between strains. For some reporter fusions, DNA fragments were PCR amplified using Q5® Hot Start High-Fidelity 2X Master Mix (NEB) and oligonucleotides described in Table S1. The wild-type and enhancer mutant *manY′-′lacZ* translational fusions (Fig. 4D) were generated using single-stranded oligos (Table S1) containing 5’ homology to P_BAD_ and 3’ homology to *lacZ.* Products were transformed into strains and λ-red homologous recombination was used to integrate fragments on the chromosome under the control of an arabinose-inducible P_BAD_ promoter.

A DNA fragment containing the *rpsA* gene, encoding r-protein S1, was PCR amplified from MG1655 genomic DNA using OSA632/OSA633 primer pair. The pET28 backbone was PCR amplified with OSA636/OSA637 primer pair using pET28-Scy-C as a template (a gift from the Nair lab, University of Illinois at Urbana). The fragments were assembled into pET28-*rpsA* plasmid using the NEBuilder® HiFi DNA Assembly Master Mix (following manufacturer’s instructions). Primer pair OSA779/OSA780 was used to PCR amplify pET28-*rpsA* and introduce mutations into the *rpsA* coding region to yield a construct encoding mS1 protein (Y205A, F208A, H219A, Fig. 6B, C). The amplified PCR product was circularized using the NEBuilder® HiFi DNA Assembly Master Mix to generate pET28-*mrpsA* (Table S2). Wild-type and mutated *rpsA* under the control of weak and or strong SD sequences were PCR amplified from pET28 constructs using OSA774/OSA776 (for strong SD) and OSA775/OSA776 (weak SD) primer pairs. Assembly of these PCR fragments with the pWKS30 backbone (amplified with the primer pair OSA783/OSA784) were performed using the NEBuilder® HiFi DNA Assembly Master Mix following the manufacturer’s instructions.

### Media and regents

Unless otherwise stated, bacteria were cultured in LB broth or on LB agar plates at 37°C. Cultures used for β-Galactosidase assays were grown in TB medium. For induction of P_Lac_ promoters, IPTG was used at a concentration of 0.1 mM. To induce P_BAD_ promoters, L-arabinose was used at concentrations of 0.001% for solid media, and 0.002% for liquid media. Antibiotics were used at the following concentrations: 100 µg/ml ampicillin, 25 µg/ml chloramphenicol, and 25 µg/ml kanamycin.

### β-Galactosidase assays

For most β-galactosidase assays, reporter strains were grown overnight in TB medium and subcultured 1:100 to fresh medium containing Amp and 0.002% L-arabinose. Cultures were grown at 37°C with shaking to OD_600_ ∼0.2, and where relevant, 0.1 mM IPTG (final concentration) was added to induce expression of SgrS. Cells were grown for another hour to OD_600_ ∼0.5. β-Galactosidase assays were performed on these cells according to the previously published protocol (Miller, 1972).

For assays conducted on strains in Fig. 8, culture conditions were slightly different. Strains with *manY′-′lacZ, thrS′-′lacZ*, and *sodB′-′lacZ* fusions harboring WT S1 or mS1 plasmids were grown overnight in TB medium and subcultured 1:100 to a fresh medium containing Amp. Cultures were grown at 37°C with shaking to OD_600_ ∼0.4, and 0.002% L-arabinose (final concentration) was used to induce expression of reporter fusions. Cells were grown for another 20 minutes and 0.1 mM IPTG (final concentration) was added to induce S1 or mS1 production. Cells were grown for an additional 10 min before β-Galactosidase assays were performed. SgrS was induced by adding 0.5% αMG (final concentration) to the media.

### In vitro transcription

For in vitro transcription, template DNA was generated by PCR using gene specific oligonucleotides with a T7 promoter sequence at the 5’ end of the forward primer. The following oligonucleotides were used to generate templates for RNA footprinting and gel shift assays: OSA753/OSA754 and OJH218/OJH169 to generate *manY* and SgrS template DNA. The oligonucleotide pair OSA734/OSA735 was used to PCR amplify *thrS* template from MG1655 genomic DNA. Transcription of these DNA templates was performed using the MEGAscript T7 kit (Ambion) following the manufacturer’s instructions.

### Purification of Hfq

Hfq-His_6_ protein was purified following a previously published protocol (Maki *et al.*, 2008). BL21(DE3) cells harboring pET21b-Hfq-His_6_ were cultured in 400 ml LB medium. At OD_600_ ∼0.3, 1 mM IPTG (final concentration) was added to the culture and incubation was continued for 2 hrs. The cells were washed with STE buffer (100 mM NaCl; 10 mM Tris·HCl, pH 8.0; 1 mM EDTA) and resuspended in 10 ml Equilibration buffer (50 mM Na_2_HPO_4_–NaH_2_PO_4_, 300 mM NaCl, and 10 mM imidazole). The suspension was treated with 25 mg lysozyme, incubated on ice for 10 minutes and sonicated. The supernatant was collected after centrifugation at 16,000 × g for 10 min at 4°C followed by incubation at 80°C for 10 minutes. The sample was centrifuged again, at 16,000 × g for 10 min at 4°C. The supernatant was fractionated using a Ni^2+^NTA agarose column following the manufacturer’s instructions (Roche) and checked by SDS-PAGE electrophoresis. The fractions containing Hfq were pooled, dialyzed, and stored in a storage buffer (20 mM Tris·HCl pH 8.0, 0.1 M KCl; 5 mM MgCl_2_, 50% glycerol, 0.1% Tween 20, and 1 mM DTT) at −20°C.

### Purification of S1

S1 was purified following a published protocol with modifications (Sukhodolets & Garges, 2003). *E. coli* BL21 (DE3) cells with the pET28-*rpsA* vector was grown to late exponential phase. Around OD of 0.6-0.8, the expression of the protein was induced with IPTG (1mM, final concentration) and the incubation was continued for 4 hr at 37 °C. At this point, cells were harvested by centrifugation and the cell pellet was resuspended in 30 mL of extraction buffer (1X PBS, 0.5 M NaCl, pH 7.2) and lysed in a French press. The supernatant was collected after centrifugation at 16,000 × g for 10 min at 4°C. The supernatant was fractionated using a Hi-Trap Ni^2+^ column (GE Healthcare) following the manufacturer’s instructions and checked by SDS-PAGE electrophoresis. The fractions containing S1 were dialyzed overnight in TGED buffer (10 mM Tris-HCl pH 8, 5% glycerol, 0.1 mM EDTA, and DTT 0.015 mg/mL) and loaded on a mono-Q column (GE Healthcare). The column was washed with TGED buffer and protein was eluted with a linear gradient of TGED buffer with NaCl (0.1 M to 1 M). The fractions containing the S1 protein were pooled, dialyzed, and concentrated using Centricon 10 concentrators (Millipore-Sigma). The poly-histidine tag was removed using the Thrombin cleavage kit (Millipore-Sigma) following manufacturer’s instructions. The cleaved proteins were further resolved in a Superdex 200 column with TGED buffer containing 0.1 M NaCl. The column fractions containing S1 were concentrated using Centricon 10 concentrator (Millipore-Sigma), mixed with an equal volume of 100% glycerol and stored at −20 °C.

### Footprinting assays

RNA footprinting reactions were performed as described previously (Desnoyers *et al.*, 2009). In brief, 0.1 pmol of 5’-end labeled *manY* mRNA was incubated with unlabeled SgrS and Hfq, and incubated at 37 °C for 10 minutes in structure buffer (Ambion) containing 1 ng of yeast RNA (Ambion). Lead acetate (Sigma) was added to perform the cleavage reaction (2.5 µM, final concentration) and incubated at 37 °C for two minutes. At this point, reactions were stopped by adding 12 µL of loading buffer II (Ambion). *manY* mRNA transcripts were incubated at 90 °C for 5 minutes in alkaline buffer (Ambion) to generate the alkaline ladder. The samples were resolved on an 8 % polyacrylamide-urea gel. For ribosome footprints, 0.1 pmol of 5’-end labeled *manY* and *thrS* mRNAs were incubated at 37 °C for 30 minutes in structure buffer (Ambion) containing 1 ng of yeast RNA (Ambion), in the presence or absence of 10 pmol *E. coli* 70S ribosome (NEB). Lead acetate mediated cleavage was performed as described above. Reactions were stopped with 12 µL of loading buffer II (Ambion). The alkaline hydrolysis ladder was generated by incubating the end-labeled transcript at 95 °C for 5 min in alkaline buffer (Ambion). RNase T1 was used for 5 min at 37 °C to generate the G ladder. The samples were resolved in an 8 % polyacrylamide-urea gel.

The DEPC footprint experiment was performed following a published protocol with some modifications (Boni *et al.*, 2001). In brief, 5 pmol of *in vitro* transcribed *manY* transcript was incubated with WT or mS1 protein (3 µM, final concentration) at 37 °C for 10 minutes in binding buffer (20 mM Tris-HCl pH 7.6, 10 mM MgCl_2_, 100 mM NH_4_Cl, and 0.5 mM DTT). 1 µL DEPC (Sigma, undiluted) was added to the reaction mixture and incubated at room temperature with gentle shaking. Reactions were stopped with 40 µL stop solution (0.4 M Na-acetate pH 5.2, and 20 mM EDTA). Modified *manY* transcripts were phenol extracted and precipitated with ethanol. Using a reverse primer (OSA754), primer extension was performed with α^32^P dATP (Perkin-Elmer) in the reaction mixture. Sanger ladders were generated using the Sequenase Version 2.0 DNA Sequencing Kit (Applied Biosystems).

### Electrophoretic mobility shift assay

RNA-RNA and RNA-protein gel electrophoretic mobility shift assays were performed using 0.01 pmol of P^32^-labeled *manY* RNA and the indicated amounts of SgrS in binding buffer (10mM Tris-HCl pH 8.0, 0.5 mM DTT, 0.5 mM MgCl_2_, 10mM KCl, 5mM Na_2_HPO_4_–NaH_2_PO_4_ pH 8.0). The mixture was incubated at 37 °C for 30 minutes, and non-denaturing loading buffer (50% glycerol and 0.1% bromophenol blue) was added. The samples were resolved on a 4.6 % native polyacrylamide gel for 1.5 hours at 10 mA. The fraction of *manY* RNA bound was determined using Fluorescent Image Analyzer FLA-3000 (FUJIFILM) to quantify the band intensities. The data were fit into Sigmaplot software to obtain the *K*_D_ value. For S1-RNA gel shift assays, 0.01 pmol of P^32^-labeled RNA and the indicated amounts of S1 (or mS1) protein TGED buffer with 0.1 M NaCl. The reaction mixture was incubated and resolved in a native gel as described above.

Hfq-sRNA-target mRNA gel mobility shift assays were performed using 0.01 pmol of P^32^-labeled *manY* RNA and the indicated amounts of SgrS and Hfq in binding buffer. The mixtures were incubated at 37°C for 30 minutes. Hfq was removed from the reaction by phenol extraction. A non-denaturing loading buffer (50% glycerol and 0.1% bromophenol blue) was added to the aqueous phase and the samples were resolved on a 4.6% native polyacrylamide gel for 1.5 hours at 10 mA. The fraction of *manY* RNA bound was determined using Fluorescent Image Analyzer FLA-3000 (FUJIFILM) to quantitate the intensities of the bands. The data were fit into GraphPad Prism7 software to obtain the *K*_D_ value.

### In vitro translation

In vitro transcribed 3X-FLAG tagged *manY* transcripts – wild-type or Δenh (1 pmol) were translated in vitro using the PURExpress translation kit (NEB). Translation reactions were stopped by adding Laemmli sample buffer (Bio-Rad). The samples were resolved in a NuPAGE gradient gel (Life Technologies) and transferred to Immobilon-P^SQ^ membrane (Millipore-Sigma). The membranes were incubated in a blocking buffer with anti-FLAG (Sigma) and anti-GFP (Thermo Scientific) antibodies. The signals were visualized with ECL western blotting substrate (Thermo Scientific).

## Acknowledgements

We thank members of the Slauch laboratory, and the present and former members of the Vanderpool laboratory for productive discussions and critical feedback. We are grateful to Marinos Kalafatis and Maxim Sokholdolets (Lamar University) for advice on S1 purification. This work was supported by the National Institutes of Health (Grants GM092830 and GM112659) and a University of Illinois Department of Microbiology graduate fellowship (provided to M.S.A.).

## Author contributions

C.K.V. and M.S.A. conceived and designed this study; M.S.A. performed the research; C.K.V. and M.S.A. analyzed the data and prepared the manuscript.

## Data Availability Statement

The data that support the findings of this study are available from the corresponding author upon reasonable request.

## Abbreviated Summary

The *manY* mRNA folds into a structure that requires a specific interaction with ribosomal protein S1, which unfolds the structure and promotes ribosome binding and translation initiation. S1 binding to *manY* mRNA thus enhances *manY* translation. SgrS, a small RNA regulator in *Escherichia coli*, forms an RNA-RNA duplex with *manY* mRNA, which interferes with S1 binding. This allows SgrS to inhibit translation of *manY* mRNA.

