## Supplemental Tables 1 and 2 for "Translation inhibition from a distance: the small RNA SgrS interferes with a ribosomal protein S1-dependent enhancer"

Carin K. Vanderpool, Ph.D.

(t) 217-333-7033

(f) 217-244-6697

Table S1. Oligonucleotides used in this study

| Oligo | Description | Sequence 5´-3´ |
|---|---|---|
| manYUTR | Single-stranded oligo for recombineering *manY* 5´ UTR into PM1205 | TCGCAACTCTCTACTGTTTCTCCATCGTATTGTGTTGATTATCACTCAGTTTTCACACTTAAGTCTTACGTAAACAGGAGAAGTACAATGGTCGTTTTACAACGTCGTGACTGGG |
| mut1 | Single-stranded oligo for recombineering *mut1 manY* 5´ UTR into PM1205 | TCGCAACTCTCTACTGTTTCTCCATCGTATTGTGTTGATTATCACTCAGGGTTCACACTTAAGTCTTACGTAAACAGGAGAAGTACAATGGTCGTTTTACAACGTCGTGACTGGG |
| mut2 | Single-stranded oligo for recombineering *mut2 manY* 5´ UTR into PM1205 | TCGCAACTCTCTACTGTTTCTCCATCGTATTGTGTTGATTATCACTCAGTTGGCACACTTAAGTCTTACGTAAACAGGAGAAGTACAATGGTCGTTTTACAACGTCGTGACTGGG |
| mut3 | Single-stranded oligo for recombineering *mut3 manY* 5´ UTR into PM1205 | TCGCAACTCTCTACTGTTTCTCCATCGTATTGTGTTGATTATCACTCAGGGGGCACACTTAAGTCTTACGTAAACAGGAGAAGTACAATGGTCGTTTTACAACGTCGTGACTGGG |
| mut4 | Single-stranded oligo for recombineering *mut4 manY* 5´ UTR into PM1205 | TCGCAACTCTCTACTGTTTCTCCATCGTATTGTGTTGATTATCACTCAGTTTTCACACGGAAGTCTTACGTAAACAGGAGAAGTACAATGGTCGTTTTACAACGTCGTGACTGGG |
| mut5 | Single-stranded oligo for recombineering *mut5 manY* 5´ UTR into PM1205 | TCGCAACTCTCTACTGTTTCTCCATCGTATTGTGTTGATTATCACTCAGAAAACACACTTAAGTCTTACGTAAACAGGAGAAGTACAATGGTCGTTTTACAACGTCGTGACTGGG |
| mut6 | Single-stranded oligo for recombineering *mut6 manY* 5´ UTR into PM1205 | TCGCAACTCTCTACTGTTTCTCCATCGTATTGTGTTGATTATCACTCATTTTTCACACTTAAGTCTTACGTAAACAGGAGAAGTACAATGGTCGTTTTACAACGTCGTGACTGGG |
| mut7 | Single-stranded oligo for recombineering *mut7 manY* 5´ UTR into PM1205 | TCGCAACTCTCTACTGTTTCTCCATCGTATTGTGTTGATTATCACTCAATTTTCACACTTAAGTCTTACGTAAACAGGAGAAGTACAATGGTCGTTTTACAACGTCGTGACTGGG |
| mut9 | Single-stranded oligo for recombineering *mut9 manY* 5´ UTR into PM1205 | TCGCAACTCTCTACTGTTTCTCCATCGTATTGTGTTGATTATCACTCAGTTTTTACACTTAAGTCTTACGTAAACAGGAGAAGTACAATGGTCGTTTTACAACGTCGTGACTGGG |
| mut10 | Single-stranded oligo for recombineering *mut10 manY* 5´ UTR into PM1205 | TCGCAACTCTCTACTGTTTCTCCATCGTATTGTGTTGATTATCACTCAGTTTTCGCACTTAAGTCTTACGTAAACAGGAGAAGTACAATGGTCGTTTTACAACGTCGTGACTGGG |
| mut14 | Single-stranded oligo for recombineering *mut14 manY* 5´ UTR into PM1205 | TCGCAACTCTCTACTGTTTCTCCATCGTATTGTGTTGATTATCACTCAGTTTTCACGCTTAAGTCTTACGTAAACAGGAGAAGTACAATGGTCGTTTTACAACGTCGTGACTGGG |
| mut15 | Single-stranded oligo for recombineering *mut14 manY* 5´ UTR into PM1205 | TCGCAACTCTCTACTGTTTCTCCATCGTATTGTGTTGATTATCACTCAGTTTTCACTCTTAAGTCTTACGTAAACAGGAGAAGTACAATGGTCGTTTTACAACGTCGTGACTGGG |
| OSA628 | T7 forward primer for *in vitro* transcription of *manY* (-129 relative to the *manY* start codon) | TAATACGACTCACTATAGGGAGTCCGTAAGGTTTCCACCGAT |
| OSA629 | Reverse primer for *in vitro* transcription of *manY* (+74) | TCGAGGATTGATCCCATACCTGCGATA |
| manY46UTR | Single-stranded oligo for recombineering *manY* 5´ UTR (46nt) into PM1205 | TCGCAACTCTCTACTGTTTCTCCATCACTCAGTTTTCACACTTAAGTCTTACGTAAACAGGAGAAGTACAATGGTCGTTTTACAACGTCGTGACTGGG |

| | | |
|---|---|---|
| Δenh | Single-stranded oligo for recombineering *manY* 5′ UTR (Δenh) into PM1205 | TCGCAACTCTCTACTGTTTCTCCATTTAAGTCTTACGTAAACAGGAGAAGTACAATGGTCGTTTTACAACGTCGTGACTGGG |
| mvENH | Single-stranded oligo for recombineering *manY* 5′ UTR (ATGenh) into PM1205 | TCGCAACTCTCTACTGTTTCTCCATTTAAGTCTTACGTAAACAGGAGAAGTACAATGCACTCAGTTTTCACACTTGTCGTTTTACAACGTCGTGACTGGG |
| OSA003 | T7 forward primer for *in vitro* translation of *manY* (-62 relative to the *manY* start codon) | TAATACGACTCACTATAGGGCGTATTGTGTTGATTATCACTCAGTTTTC |
| OSA823 | Reverse primer for *in vitro* translation of *manY*-3XFLAG (+351) | TATTTGATGCCTCTAGAGTCTTACTTATCGTCATCGTCTTTGTAATCAATATCATGATCCTTGTAGTCTCCGTCGTGGTCCTTATAGTCGGTAATAGTACGAACGATGATGGTCAGTACCT |
| OSA321 | T7 forward primer for in vitro translation of *manY* (-62 relative to the *manY* start codon, Δenh) | TAATACGACTCACTATAGGGCGTATTGTGTTGATTATTTAAGTCTTACGTAAACAGGAGAAGTAC |
| Y UTR lacZ SD | Single-stranded oligo for recombineering *manY* 5′ UTR with *lacZ* SD into PM1205 | TCGCAACTCTCTACTGTTTCTCCATCGTATTGTGTTGATTATCACTCAGTTTTCACACTTAAGTCTTACGTAAACAGGAAACAGCTATGGTCGTTTTACAACGTCGTGACTGGG |
| lacZ mut3 | Single-stranded oligo for recombineering *manY* 5′ UTR (with *lacZ* SD and *mut3* enhancer) into PM1205 | TCGCAACTCTCTACTGTTTCTCCATCGTATTGTGTTGATTATCACTCAGGGGGCACACTTAAGTCTTACGTAAACAGGAAACAGCTATGGTCGTTTTACAACGTCGTGACTGGG |
| lacZ mut5 | Single-stranded oligo for recombineering *manY* 5′ UTR (with *lacZ* SD and *mut5* enhancer) into PM1205 | TCGCAACTCTCTACTGTTTCTCCATCGTATTGTGTTGATTATCACTCAGAAAACACACTTAAGTCTTACGTAAACAGGAAACAGCTATGGTCGTTTTACAACGTCGTGACTGGG |
| flip-enh | Single-stranded oligo for recombineering *manY* 5′ UTR (with the flipped enhancer) into PM1205 | TCGCAACTCTCTACTGTTTCTCCATCTATTAGTTGTGTTATGCACTCAGTTTTCACACTTCAAATGCATTCTGAAAGGAGAAGTACAATGGTCGTTTTACAACGTCGTGACTGGG |
| sSD | Single-stranded oligo for recombineering *manY* 5′ UTR (with a strong SD) into PM1205 | TCGCAACTCTCTACTGTTTCTCCATCGTATTGTGTTGATTATCACTCAGTTTTCACACTTAAGTCTTACGTAAACAGGAGGAGTACAATGGTCGTTTTACAACGTCGTGACTGGG |
| wSD | Single-stranded oligo for recombineering *manY* 5′ UTR (with a weak SD) into PM1205 | TCGCAACTCTCTACTGTTTCTCCATCGTATTGTGTTGATTATCACTCAGTTTTCACACTTAAGTCTTACGTAAATTGGAGTAGTACAATGGTCGTTTTACAACGTCGTGACTGGG |
| sSD+mut3 | Single-stranded oligo for recombineering *manY* 5′ UTR (with a strong SD and the *mut3* enhancer) into PM1205 | TCGCAACTCTCTACTGTTTCTCCATCGTATTGTGTTGATTATCACTCAGGGGGCACACTTAAGTCTTACGTAAACAGGAGGAGTACAATGGTCGTTTTACAACGTCGTGACTGGG |

| | | |
|---|---|---|
| wSD+mut3 | Single-stranded oligo for recombineering *manY* 5′ UTR (with a weak SD and the *mut3* enhancer) into PM1205 | TCGCAACTCTCTACTGTTTCTCCATCGTATTGTGTTGATTATCACTCAGGGGGCACACTTAAGTCTTACGTAAATTGGAGTAGTACAATGGTCGTTTTACAACGTCGTGACTGGG |
| WT thrS | Single-stranded oligo for recombineering *thrS* 5′ UTR (-66 to -1, relative to the start codon) into PM1205 | TCGCAACTCTCTACTGTTTCTCCATTAATAAACAAATTTTTCTTTGTATGTGATCTTTCGTGTGGGTCACCACTGCAAATAAGGATATAAAATGGTCGTTTTACAACGTCGTGACTGGG |
| A-thrS | Single-stranded oligo for recombineering *thrS* 5′ UTR with *A-thrS* (-66 to -1, relative to the start codon) into PM1205 | TCGCAACTCTCTACTGTTTCTCCATTAATAAACAAAAAAAACTTTGTATGTGATCTTTCGTGTGGGTCACCACTGCAAATAAGGATATAAAatgGTCGTTTTACAACGTCGTGACTGGG |
| G-thrS | Single-stranded oligo for recombineering *thrS* 5′ UTR with *G-thrS* (-66 to -1, relative to the start codon) into PM1205 | TCGCAACTCTCTACTGTTTCTCCATTAATAAACAAAGGGGGCTTTGTATGTGATCTTTCGTGTGGGTCACCACTGCAAATAAGGATATAAAATGGTCGTTTTACAACGTCGTGACTGGG |
| OSA734 | T7 forward primer for *in vitro* translation of *thrS* (-66 relative to the *thrS* start codon) | TAATACGACTCACTATAGGGGATTGCGAACCAATTTAGCATTTGTTGG |
| OSA753 | T7 forward primer for *in vitro* translation of *manY* (-98 relative to the *manY* start codon, for ribosome footprint, and reverse transcription of DEPC-modified transcripts) | TAATACGACTCACTATAGGGAAAATGATGGATCTGATCAGCAAAATCGATAAGT |
| OSA754 | Reverse primer for *in vitro* transcription of *manY* (+98 relative to the *manY* start codon, for ribosome footprint, and reverse transcription of DEPC-modified transcripts) | AGCGGACGGTGAAACTGAAATTC |
| OSA786 | Forward primer for recombineering *sodB* (+1, transcription start site) into PM1205 | ACCTGACGCTTTTTATCGCAACTCTCTACTGTTTCTCCATATACGCACAATAAGGCTATTGTACGTATGC |
| OSA787 | Reverse primer for recombineering *sodB* (+120, relative to the start codon) into PM1205 | TAACGCCAGGGTTTTCCCAGTCACGACGTTGTAAAACGACGTTCAGGTTAGTGACATAAGTCTGATGGT |
| OSA070 | T7 forward primer for *in vitro* transcription and translation of *gfp* | TAATACGACTCACTATAGGGACGTAAAGGAGGAATTCATGAGCAAAGGAGAAGAACTTTTCACTG |

| | | |
|---|---|---|
| OSA071 | Reverse primer for *in vitro* transcription and translation of *gfp* | TACGTTTATTTGTAGAGCTCATCCATGCCATGTG |
| OSA632 | Forward primer to PCR amplify *rpsA* | CTTTAAGAAGGAGATATACCATGACTGAATCTTTTGCTCAACTCTTTGAAGAG |
| OSA633 | Reverse primer to PCR amplify *rpsA* | CTGCTGCCGCGCGGCACCAGCTCGCCTTTAGCTGCTTTGAAAGCTT |
| OSA636 | Forward primer for pET28 | TTGATCTTTTCTACGGGGTCTGACGCT |
| OSA637 | Reverse primer for pET28 | CTTCTTGAGATCCTTTTTTTCTGCGCGT |
| OSA779 | Primer to introduce Y205A, F208A, and H219A mutations in *rpsA* | GTTGATCTGGGCGGCGTTGACGGCCTGCTGGCTATCACTGACATGGCCTGGAAACG |
| OSA780 | Primer to introduce Y205A, F208A, and H219A mutations in *rpsA* | TCAACGCCGCCCAGATCAACGGCTGCACCGGCGTCAGTGAGGTTCTTAACGATACCTTTAACTTCC |
| OSA774 | Forward primer with a strong SD to PCR amplify *rpsA* and *mrpsA* (to construct pWKS30-SS1) | TTGTGAGCGGATAACAATTTCACACATTGGAGGAACAATATGACTGAATCTTTTGCTCAACTCTTTGAAG |
| OSA775 | Forward primer with a weak SD to PCR amplify *rpsA* and *mrpsA* (to construct pWKS30-WS1) | TTGTGAGCGGATAACAATTTCACACATTCCAGGAACAATATGACTGAATCTTTTGCTCAACTCTTTGAAG |
| OSA776 | Reverse primer to PCR amplify *rpsA* and *mrpsA* (to construct pWKS30-WS1) | GTTGTAAAACGACGGCCAGTATCCGGATATAGTTCCTCCTTTCAGCA |
| OSA783 | Forward primer for pWKS30 (for Gibson assembly) | ACTGGCCGTCGTTTTACAACG |
| OSA784 | Reverse primer for pWKS30 (for Gibson assembly) | AAATTGTTATCCGCTCACAATTCCACACA |
| OJH218 | T7 forward primer for *in vitro* transcription of SgrS | TAATACGACTCACTATAGGGGATGAAGCAAGGGGGTGCCCCATG |
| OJH169 | Reverse primer for *in vitro* transcription of SgrS | AAAAAAAACCAGCAGGTATAATCTGCTGGCGGG |

Table S2. Strains and plasmids used in this study

| Plasmid | Genotype | Source |
|---|---|---|
| pBRCS12 | pHDB3, $P_{LlacO}$, vector control | (Wadler & Vanderpool, 2009) |
| pLCV1 | pHDB3 $P_{LlacO}$-*sgrS*$_{K12}$ | (Vanderpool & Gottesman, 2004) |
| pBRCS22 | pHDB3 $P_{LlacO}$-*sgrS*$_{St}$ | (Wadler & Vanderpool, 2009) |
| pWKS30 | $P_{LlacO}$, vector control | (Wang & Kushner, 1991) |
| pWKS30-SmS1 | $P_{Lac}$-strong SD-*mrpsA* | This study |
| pWKS30-WmS1 | $P_{Lac}$-weak SD-*mrpsA* | This study |
| pWKS30-WS1 | $P_{Lac}$-weak SD-WT *rpsA* | This study |
| pWKS30-SS1 | $P_{Lac}$-strong SD-WT *rpsA* | This study |
| pET28-rpsA | T7 promoter –WT *rpsA*-thrombin site- $(His)_{12}$ | This study |
| pET28-mrpsA | T7 promoter –*mrpsA*-thrombin site- $(His)_{12}$ | This study |
| **Strain** | **Genotype** | **Source** |
| PM1205 | *lacI′:: $P_{BAD}$-cat-sacB-lacZ, miniλtet$^R$, ΔaraBAD araC+, mal::lacI$^q$* | (Mandin & Gottesman, 2009) |
| SA1747 | $P_{BAD}$-*manY* UTR-ATG*'-'lacZ, ΔaraBAD araC+, mal::lacI$^q$* | This study |
| SA1735 | $P_{BAD}$-*mut1*-ATG*'-'lacZ, ΔaraBAD araC+, mal::lacI$^q$* | This study |
| SA1736 | $P_{BAD}$-*mut2*-ATG*'-'lacZ, ΔaraBAD araC+, mal::lacI$^q$* | This study |
| SA1737 | $P_{BAD}$-*mut3*-ATG*'-'lacZ, ΔaraBAD araC+, mal::lacI$^q$* | This study |
| SA1738 | $P_{BAD}$-*mut4*-ATG*'-'lacZ, ΔaraBAD araC+, mal::lacI$^q$* | This study |
| SA1739 | $P_{BAD}$-*mut5*-ATG*'-'lacZ, ΔaraBAD araC+, mal::lacI$^q$* | This study |
| SA1748 | $P_{BAD}$-*mut6*-ATG*'-'lacZ, ΔaraBAD araC+, mal::lacI$^q$* | This study |
| SA1749 | $P_{BAD}$-*mut7*-ATG*'-'lacZ, ΔaraBAD araC+, mal::lacI$^q$* | This study |
| SA1801 | $P_{BAD}$-*WT thrS* UTR*'-'lacZ, ΔaraBAD araC+, mal::lacI$^q$* | This study |
| SA1811 | $P_{BAD}$-*G-thrS'-'lacZ, ΔaraBAD araC+, mal::lacI$^q$* | This study |
| SA1815 | $P_{BAD}$-*A-thrS'-'lacZ, ΔaraBAD araC+, mal::lacI$^q$* | This study |
| SA1516 | $P_{BAD}$-*manY* UTR(*Δenh*)*'-'*ATG-*lacZ, ΔaraBAD araC+, mal::lacI$^q$* | This study |
| SA1517 | $P_{BAD}$-*manY* UTR(*Δenh*)*'-'*ATG-*enh*(in frame)-*lacZ, ΔaraBAD araC+, mal::lacI$^q$* | This study |
| SA1806 | $P_{BAD}$-flip-enh *manY* UTR*'-'lacZ, ΔaraBAD araC+, mal::lacI$^q$* | This study |
| SA1807 | $P_{BAD}$-*manY* UTR with a strong SD and *mut3* enhancer*'-'lacZ, ΔaraBAD araC+, mal::lacI$^q$* | This study |

| | | |
|---|---|---|
| SA1808 | $P_{BAD}$-*manY* UTR with a *lacZ* SD and *mut5'-'lacZ, ΔaraBAD araC+, mal::lacI$^q$* | This study |
| SA1809 | $P_{BAD}$-*manY* UTR with a weak SD*'-'lacZ, ΔaraBAD araC+, mal::lacI$^q$* | This study |
| SA1810 | $P_{BAD}$-*manY* UTR with a strong SD*'-'lacZ, ΔaraBAD araC+, mal::lacI$^q$* | This study |
| SA1916 | $P_{BAD}$-*mut9* ATG*´-'lacZ, ΔaraBAD araC+, mal::lacI$^q$* | This study |
| SA1917 | $P_{BAD}$-*mut10* ATG*'-'lacZ, ΔaraBAD araC+, mal::lacI$^q$* | This study |
| SA1918 | $P_{BAD}$-*mut14* ATG*'-'lacZ, ΔaraBAD araC+, mal::lacI$^q$* | This study |
| SA1919 | $P_{BAD}$-*mut15* ATG*'-'lacZ, ΔaraBAD araC+, mal::lacI$^q$* | This study |
| SA1907 | $P_{BAD}$-*manY* UTR-ATG*'-'lacZ, hfq::kan, ΔaraBAD araC+, mal::lacI$^q$* | This study |
| SA1902 | $P_{BAD}$-*sodB'-'lacZ, ΔaraBAD araC+, mal::lacI$^q$* | This study |
